# Sodium channel endocytosis drives axon initial segment plasticity

**DOI:** 10.1101/2022.11.09.515770

**Authors:** Amélie Fréal, Nora Jamann, Jolijn Ten Bos, Jacqueline Jansen, Naomi Petersen, Thijmen Ligthart, Casper C. Hoogenraad, Maarten H. P. Kole

**Affiliations:** Axonal Signaling Group, Netherlands Institute for Neurosciences (NIN), Royal Netherlands Academy for Arts and Sciences (KNAW); Amsterdam, The Netherlands; Cell Biology, Neurobiology and Biophysics, Department of Biology, Faculty of Science, Utrecht University; Utrecht, The Netherlands; Department of Neuroscience, Genentech, Inc; South San Francisco, United States

## Abstract

Activity-dependent plasticity of the axon initial segment (AIS) endows neurons with the ability to adapt action potential output to changes in network activity. Action potential initiation at the AIS highly depends on the clustering of voltage-gated sodium channels, however the molecular mechanisms regulating their plasticity remain largely unknown. Here, we used novel genetic tools to endogenously label sodium channels and their scaffolding protein, to reveal their nanoscale organization and longitudinally image AIS plasticity in hippocampal neurons, in slices and primary cultures. We find that induction of NMDA receptor-mediated long-term synaptic depression is linked to a rapid and local endocytosis of sodium channels from the distal AIS. These data reveal a novel fundamental mechanism for rapid activity-dependent AIS reorganization sharing conserved features with synaptic plasticity.

## Results

The axon initial segment (AIS) is a critical domain of neurons which maintains neuronal polarity and functionally acts as the final synaptic integration site for action potential (AP) generation (Kole & Stuart, 2012; Leterrier, 2018). The threshold and upstroke dynamics of neuronal APs rely, in part, on the specific distribution, density, and isoforms of voltage-gated sodium (Na_V_) channels expressed along the AIS. The AIS localization of Na_V_1.2 and Na_V_1.6 channels, the two main isoforms of Na_V_ channels in excitatory neurons is mediated by the interaction with the AIS scaffold protein Ankyrin G (AnkG) (Garrido et al., 2003; Lemaillet, Walker, & Lambert, 2003). Over the last decade emerging evidence showed that AIS morphology changes with activity in a brain region- and cell type-dependent manner. *In vivo* experiments demonstrated that changes in sensory input streams induce bidirectional changes of AIS length with temporal scales of one hour to several weeks, mediating a homeostatic plasticity of action potential output (Galliano et al., 2021; Jamann et al., 2021; Kuba, Oichi, & Ohmori, 2010). The degree of AIS plasticity varies from subtle changes in ion channel isoform expression (Kuba, Yamada, Ishiguro, & Adachi, 2015), relocation of the AIS along the axonal membrane (Grubb & Burrone, 2010), changes in AIS length (Kuba et al., 2010) or to a complete proteolysis of AIS proteins during excitotoxicity (Benned-Jensen et al., 2016; Schafer et al., 2009; Zhao et al., 2020). However, in contrast to the detailed understanding of synaptic plasticity mechanisms (Roth, Zhang, & Huganir, 2017), the precise mechanisms of AIS plasticity are far from being understood. A comprehensive molecular model for how AIS plasticity is induced and expressed is still lacking. Molecular players implicated in activity-dependent AIS relocation include L-type voltage-gated calcium (Ca^2+^) channels, calcineurin, myosin II/phospho-myosin light chain, and/or the AKT-ULK1 pathway (Berger et al., 2018; Chand, Galliano, Chesters, & Grubb, 2015; Evans, Dumitrescu, Kruijssen, Taylor, & Grubb, 2015; Evans et al., 2013; Evans, Tufo, Dumitrescu, & Grubb, 2017; Ferreira da Silva et al., 2021; Grubb & Burrone, 2010). A major limitation in our understanding of AIS plasticity mechanisms is that they mostly have been studied by comparing the AIS across large populations of differentially treated neurons. Live reporters allowing longitudinal analysis of AIS structure without affecting its organization and function are still lacking (Dumitrescu, Evans, & Grubb, 2016; Leterrier et al., 2017). Moreover, while most studies have focused on the scaffold protein AnkG, little is known about the redistribution of Na_V_ channels during plasticity. During AIS development, two main trafficking routes have been reported for membrane proteins. They are either inserted into the somatic and/or axonal membrane before diffusing laterally until they get immobilized at the AIS, like NF186 and K_V_7.3 (Ghosh, Malavasi, Sherman, & Brophy, 2020), or they are directly targeted into the AIS membrane, as has been shown for Na_V_1.6 channels (Akin, Sole, Dib-Hajj, Waxman, & Tamkun, 2015). Nevertheless, in both cases, their stable accumulation at the AIS depends on the local inhibition of their internalization by the interaction with AnkG and adaptor proteins (Eichel et al., 2022; Freal et al., 2019; Sole, Wagnon, Akin, Meisler, & Tamkun, 2019). Here, we hypothesized that, during activity-dependent AIS plasticity, analogous to synaptic plasticity, membrane channel reorganization is mediated by the dynamic trafficking of Na_V_1 channels. To resolve the molecular events underlying AIS plasticity in live neurons, we took advantage of a newly developed knock-in mouse line expressing GFP-labelled AnkG and used the ORANGE CRISPR/Cas9 system (Willems et al., 2020) to tag endogenous Na_V_1 channels, allowing the interrogation of the molecular signaling pathways during activity-induced structural plasticity of the AIS.

### NMDAR activation induces both synaptic depression and AIS plasticity

To image the AIS of individual pyramidal neurons in real time we took advantage of a recently generated AnkG-GFP knock-in mouse line and performed bilateral injections with pENN.AAV.CaMKII 0.4.Cre.SV40 into the CA1 area to obtain Cre-dependent expression of AnkG-GFP from the CaMKII promotor (**Fig. 1A**). Confocal imaging in acute hippocampal slices showed that AnkG-GFP fluorescence was detectable in the proximal axon of most pyramidal neurons along the pyramidal layer (**Fig. 1A**), and the GFP signal colocalized with the AIS markers AnkG, β34-spectrin (**fig. S1A**) and Na_V_1.6 (*n* = 10 AIS, **fig. S1B**). To induce activity-dependent plasticity we modified a previously developed approach (H. K. Lee, Kameyama, Huganir, & Bear, 1998) and induced chemical long-term synaptic depression (c-LTD) by brief exposure of slices to the N-methyl-D-aspartate (NMDA) receptor agonist NMDA (20 µM). In acute slices from AnkG-GFP mice we used an extracellular electrode to stimulate Schaffer collaterals (SC) and recorded evoked excitatory postsynaptic currents (EPSCs) from CA1 pyramidal neurons in whole-cell configuration (**Fig.1B**). NMDA was applied for 3 min and rapidly followed by wash-in of the NMDAR-selective antagonist 2-amino-5-phosphonovaltuderic acid (APV, 100 µM, 5 min, **Fig. 1B,C**). NMDA exposure caused brief action potential firing of CA1 pyramidal neurons (4.4 ± 2 min, **Fig. 1D**) followed by LTD of synaptic responses in comparison to APV exposure alone (60 min, ∼30% c-LTD vs ∼6% Control, **Fig. 1C**). Interestingly, when we simultaneously imaged AnkG-GFP fluorescence at various time points before and after NMDA application we observed a significant decrease in AIS length already 30 min after induction of c-LTD (p < 0.05, and at 60 min, 13% or ∼3.8 µm length reduction, p < 0.001, **Fig. 1E**). In imaging sessions without NMDA application no AIS shortening was observed, excluding technical confounding factors by AnkG-GFP imaging (p = 0.17, 0 vs 60 min, **Fig. 1E**). The distance of the AIS onset relative to the soma remained unchanged suggesting that the distal region of the AIS is the principal site of plasticity (**fig. S1C**). Furthermore, in separate experiments when we examined the effects of NMDA application on AIS length in slices which were not used for electrophysiological recordings, β34-spectrin staining confirmed that NMDA induced a 9% shortening of the AIS within 30 min (*n* > 200 neurons per condition**, fig. S1D**).

**Fig. 1.**
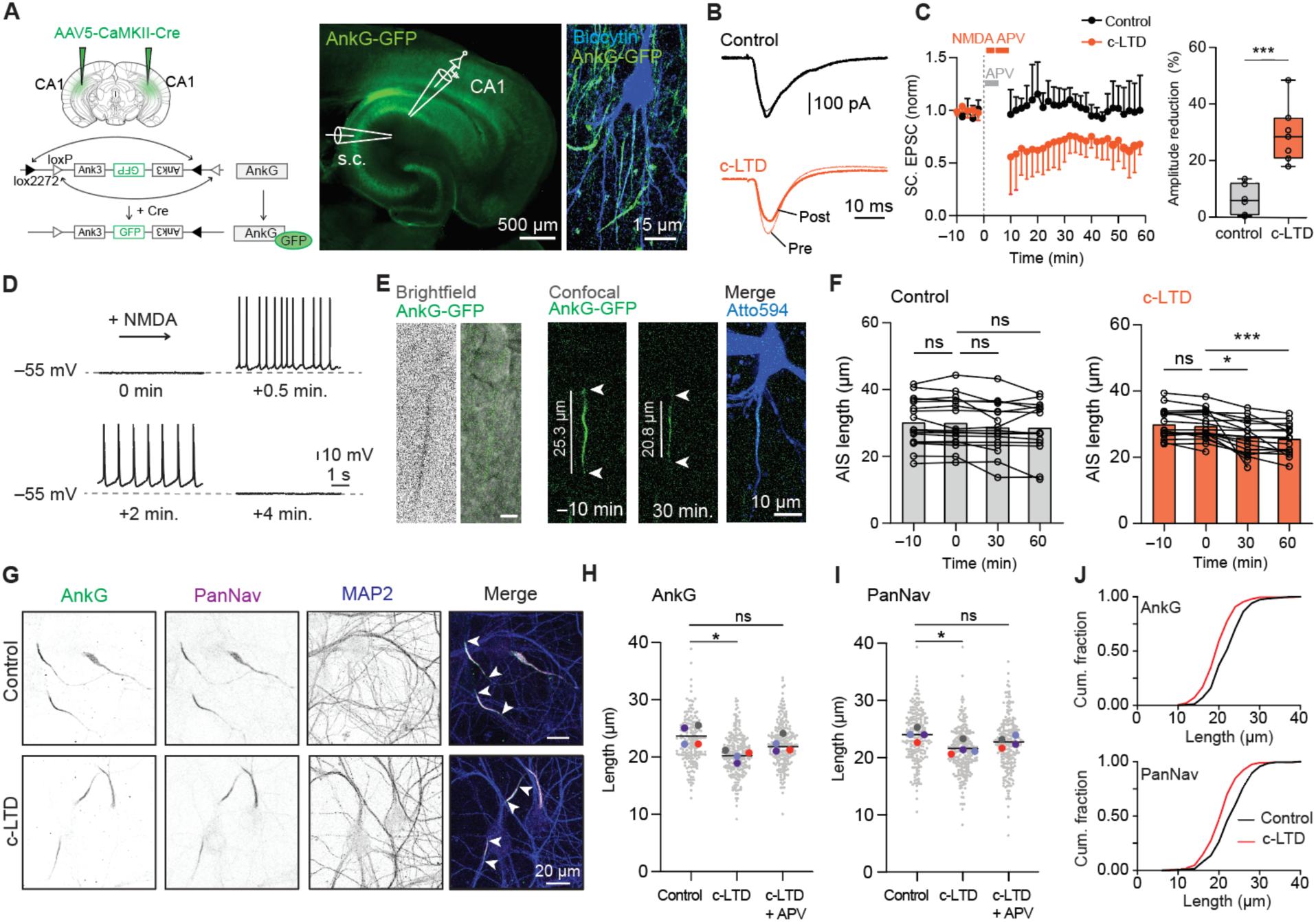
NMDAR activation induced LTD and AIS shortening in hippocampal slices and primary cultures. (**A**) AAV injection into CA1 induces expression of AnkG-GFP in CA1 pyramidal neurons, which were recorded with whole-cell patch-clamp while stimulating Schaffer collaterals (SC). Example CA1 pyramidal neuron in an acute slice filled with biocytin (blue) and immunostained against GFP (green). (**B**) Example EPSC traces before (pre) and after 60 min (post) with brief APV (control) or NMDA+APV application (c-LTD). (**C**) NMDA exposure significantly reduces EPSC amplitude (amplitude reduction (∼30% c-LTD vs ∼6% Ctrl, unpaired t-test p < 0.001). (**D**) Application of NMDA induces a ∼5 min depolarization and action potential firing. (**E**) Left, neurons bearing a GFP-positive AIS were identified with brightfield illumination. Right, confocal *z*-stacks were imaged before (–10 min) and after NMDA application (30 min). Atto594 filling confirms the overlap of AIS GFP with the neuron. AIS onset, end and length estimate indicated with white arrows. (**F**) Quantification of AIS length of recorded neurons in control (*n* = 17, 9 whole-cell, 8 imaging only) vs c-LTD (*n* = 19, 8 whole-cell, 11 imaging only) conditions. The c-LTD causes a significant AIS length reduction (Mixed effect analysis p < 0.001, Multiple comparisons: 0 vs 30 p < 0.05, 0 vs 60 p < 0.001). (**G**) Immunostaining of AnkG, PanNav and MAP2 on control hippocampal primary neurons and after induction of c-LTD (4 min 50 µM NMDA). Scale bar is 20 µm. **H-I**: Population data of AnkG (**H**) and PanNav (**I**) lengths in control, c-LTD and c-LTD + APV conditions (*N* = 4 cultures, *n* > 440 neurons per condition). Matched Friedman test with Dunn’s multiple comparisons test. AnkG, c-LTD * p = 0.027, LTD + APV p = 0.96 (ns). PanNav, c-LTD * p = 0.027, LTD + APV p = 0.15 (ns). (**J**) Plots of cumulative fractions of AnkG and PanNav lengths in control and c-LTD conditions.

Next, we addressed the effects of c-LTD induction on AIS length in cultured hippocampal neurons. We treated DIV14 primary neurons with 50 µM of NMDA (4 min) and determined the AIS length after 30 min recovery. We observed a significant reduction of AnkG (∼20%) and total Na_V_ population (PanNav) (∼10%) length respectively (**Fig. 1G-I**), which was blocked by APV (100 µM, 3 min before and during NMDA, **Fig. 1H,I**). These results are consistent with the results obtained in acute slices, and for the first time show that NMDA receptor activation causes a rapid form of AIS plasticity. A leftward shift in the cumulative distribution plots indicated that the NMDA-induced shortening triggered a global reduction in length of the AIS in all neurons rather than a strong reduction in a subpopulation (**Fig. 1J**). Together, these data show that the synaptic induction of a classic form of LTD is associated with the robust and rapid shortening of the AIS, both in hippocampal acute slices and cultured neurons.

### Loss of Nav following NMDAR activation specifically occurs in the distal AIS

The AIS of hippocampal excitatory neurons contains both sodium channel isoforms Na_V_1.2 and Na_V_1.6 (Lorincz & Nusser, 2010). To test whether NMDA-induced AIS plasticity selectively affects one or both isoforms, we first characterized their distribution along the AIS in primary cultured neurons. Staining of DIV14 hippocampal neurons revealed that 90% and 80% of AnkG-positive AISs were also positive for Na_V_1.2 and Na_V_1.6 respectively, with Na_V_1.2 expressed along the entire AIS, whereas Na_V_1.6 was predominantly found in the proximal AIS (**fig. S2A**). At DIV21 Na_V_1.6 expression became more prominent in comparison to Na_V_1.2 (**fig. S2B–D**). In DIV14 cultures, brief application of NMDA significantly reduced the length of Na_V_1.2, but not Na_V_1.6 (**fig. S3A-B**), by 14% compared to control condition in ∼75% of the experiments (*n* = 21 cultures), which was blocked by APV (**Fig. 2A,B**).

**Fig. 2.**
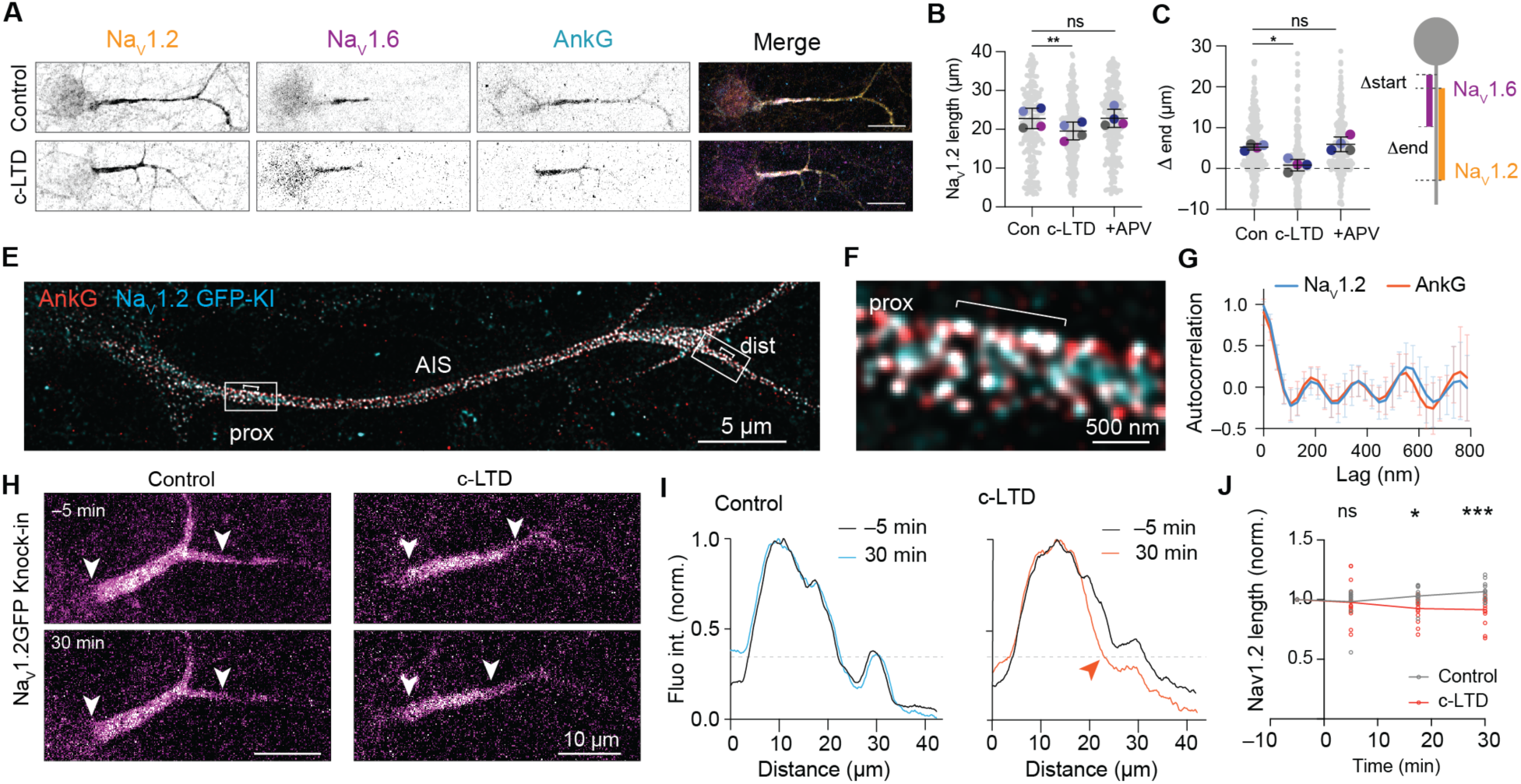
Na_V_1.2 is selectively reduced in the distal AIS during NMDAR-mediated plasticity. (**A**) Immunostaining for Na_V_1.2, Na_V_1.6 and AnkG on DIV14 in control condition and after c-LTD. Scale bars are 20 µm. (**B**) Average length of AIS Na_V_1.2 reduced by NMDA exposure and prevented by NMDA antagonist APV (c-LTD + APV). Repeated-measures One-Way ANOVA with Dunnett’s multiple comparisons test, N = 4. c-LTD, **p = 0.008, c-LTD + APV, ns p = 0.99). At least 236 neurons per condition. (**C, D**) NMDAR mediated reduction in Δ end (Na_V_1.2-Na_V_1.6). (C) Repeated-measure One-way ANOVA with Dunnett’s multiple comparisons test, N = 4 cultures. c-LTD, *p = 0.022, c-LTD+APV, ns p = 0.76. At least 260 neurons per condition, and corresponding scheme (D). (**E-G**). STED image of DIV14 Na_V_1.2-GFP knock-in neuron stained for GFP and AnkG (**E**) and zoom in the proximal AIS (**F**). Scale bar are 5 µm and 500 nm in the zoom. Autocorrelation of AnkG and Na_V_1.2 fluorescence profiles in the proximal AIS (G) *n* = 8 neurons. (**H)** Live-cell imaging of Na_V_1.2-GFP knock-in neuron 5 min before and 30 min after induction of c-LTD and control. Scales bars are 10 µm, white arrowheads point to the start and end point of the Na_V_1.2-GFP signal. (**I**) Corresponding fluorescence intensities normalized and smoothened over 1 µm. The dashed lines delineate the thresholds of end position. (**J**) Average Na_V_1.2-GFP length of control neurons (gray, *n* = 13 neurons in *N* = 2 cultures) or in c-LTD condition (red, *n* = 15 in *N* = 2 cultures) normalized to the first frame. Mixed-effects analysis with Šídák’s multiple comparisons test, 5 min, ns p > 0.99; 17.5 min, *p = 0.019; 30 min, ***p = 0.0009.

We compared the intensity and the integrated intensity of Na_V_1 isoform staining after NMDA application, normalized to the control condition in each experiment. While neither Na_V_1.6 staining mean intensity nor density changed after NMDA exposure, Na_V_1.2 staining density was reduced by 25%, with no change in the mean intensity (**fig. S3C**). These findings suggest that the NMDAR-induced shortening may reflect a local channel removal rather than a global loss of Na_V_1.2 channels. To further explore the location of NMDAR-induced changes we next measured Na_V_1.2 and Na_V_1.6 relative start and end positions to determine the location of AIS shorting (**Fig. 2C,D** and **fig. S3D**). NMDAR activation changed the relative start positions of Na_V_1.6 and Na_V_1.2 (∼2.2 µm reduction) and strongly impacted the distal Na_V_1.2 region (∼5 µm reduction), indicating that the length reduction of Na_V_1.2 reflects primarily changes at the distal end of the AIS (**Fig. 2C**).

To monitor Na_V_1.2 channel membrane dynamics in living cells without over-expression artefacts, we used the ORANGE CRISPR/Cas9 system to fluorescently tag the endogenous Na_V_1.2 channel (Willems et al., 2020). Insertion of GFP in the C-terminal part of the channel revealed a strong enrichment of Na_V_1.2 at the AIS, while lower GFP levels could be detected in the distal axon and somato-dendritic compartment, (**fig. S4)** as previously reported (Liu, Wang, Pitt, & Liu, 2022). Using STED microscopy, we observed a colocalization of AnkG with Na_V_1.2 (GFP) adopting a ring-like organization throughout the AIS (**Fig. 2E,F** and **fig. S3C**). The auto-correlation analysis revealed a highly periodic arrangement of the two proteins, both in the proximal and distal AIS (**Fig. 2G and fig. S4C-F**) with a similar peak-to-peak distance of ∼190 nm (fig. S2O), as previously observed for Na_V_1.2 and Na_V_1.6 (Liu et al., 2022). Moreover, induction of c-LTD caused a ∼10% reduction of the Na_V_1.2-GFP knock-in (**fig. S4A,B**), indicating that our tagging approach does not impair Na_V_1.2 channel behavior during plasticity (**Fig. 2A,B**).

We then performed time-lapse imaging of endogenous Na_V_1.2-GFP before, during and after c-LTD (**Fig. 2H-J**) and observed the reduction in GFP fluorescence after 30 min specifically in the distal AIS (**Fig. 2H,I**). Strikingly, the average normalized GFP length was already significantly different from the control 17 min after incubation with NMDA (∼10%, **Fig. 2J**). Altogether, our data show converging lines of evidence for a rapid activity-dependent remodeling of the AIS, selectively affecting distally localized Na_V_1.2 channels. The greater degree of plasticity in this subcellular domain implies the presence of specific mechanisms initiated by NMDAR activation and locally controlling Na_V_ isoform distribution.

### Endogenous Na_V_1.2 displays a higher mobility in the distal AIS

Next, we assessed if local differences in Na_V_1.2 behavior in basal conditions could explain its preferential loss from the AIS distal region. We first performed a detergent extraction experiment on live neurons to address the tethering of Na_V_ channels to the submembrane cytoskeleton (Winckler, Forscher, & Mellman, 1999) (**fig. S5A-C**). After extraction, we observed a significantly shortened AIS when stained for AnkG (16 ± 1.2% reduction) as well as Na_V_1.2 (26 ± 2.2%) and Na_V_1.6 (17 ± 1.6%, **fig. S5A,B**). The analysis of Na_V_1.2 and Na_V_1.6 relative expression patterns revealed a large loss of Na_V_1.2 after extraction specifically in the distal AIS (**fig. S4C**), indicating that the anchoring of this channel is less stable in this region. To test this observation with an alternative method, we performed fluorescence recovery after photobleaching (FRAP) experiments to measure the mobility of endogenously-tagged Na_V_1.2-GFP in DIV14 neurons (**Fig. 3A,B**). We bleached two regions within the AIS to be able to compare the recovery in the proximal *versus* the distal part. We measured a significantly larger recovery (∼10% average recovery after 45 min) in the distal AIS compared to the proximal region (∼7%, **Fig. 3B, C**). Comparisons of distal versus proximal recovery percentage for multiple experiments confirmed a significant ∼1.5-fold higher mobility of Na_V_1.2 in the distal AIS (**Fig. 3D**), confirming extraction experiments.

**Fig. 3.**
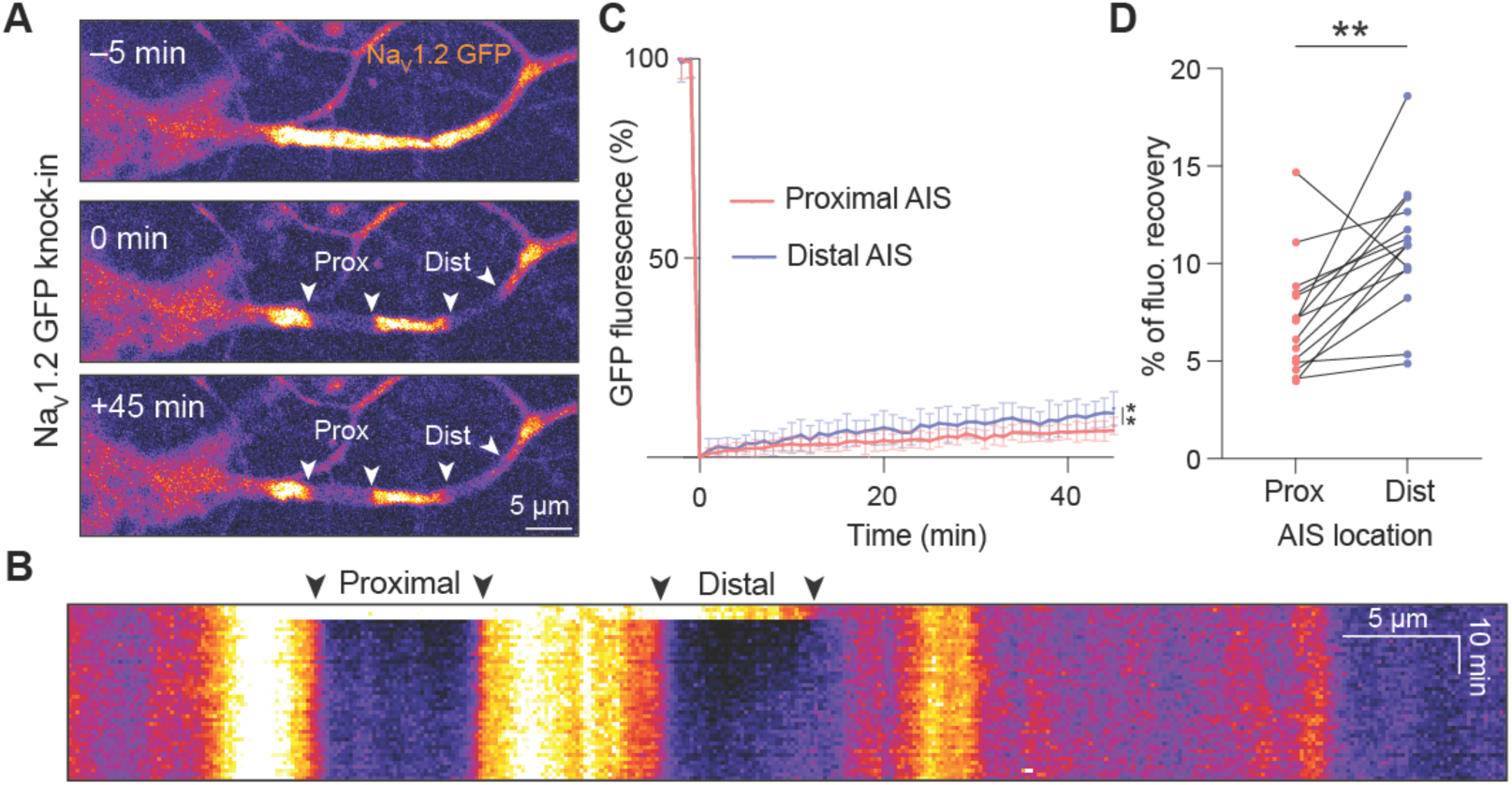
Endogenously tagged Na_V_1.2 displays higher mobility at the distal AIS. (**A**) FRAP of Na_V_1.2-GFP knock-in neurons. Still images of a Nav1.2-GFP knock-in neuron before (–5 min), and 0 min and 45 min after FRAP in the proximal and distal area (A), FRAP ROIs are indicated by white arrowheads. (**B**) Corresponding kymograph from experiment in A. (**C**) Average fluorescence recovery in the proximal and distal AIS of 15 neurons, from 2 independent experiments. 2-way ANOVA with Šídák’s multiple comparisons test, 44 min: *p = 0.02, 45 min: *p = 0.04. (**D**). Individual recovery in the proximal versus distal AIS, paired t-test **p = 0.002, *n* = 15 neurons from 2 independent experiments.

### Synaptic NMDAR-mediated depolarization triggers AIS plasticity

Although some studies reported the presence of NMDARs at axonal presynaptic release terminals (H. H. Wong, Rannio, Jones, Thomazeau, & Sjostrom, 2021), they are typically not present at the AIS (Christie & Jahr, 2009). To unravel the precise localization of NMDARs in DIV14 primary hippocampal neurons we used ORANGE to extracellularly tag the endogenous obligatory subunit GluN1 of the NMDAR with GFP (Willems et al., 2020) (**Fig. 4A,B**). We first stained for GFP without permeabilization and detected faint surface expression of GluN1 in the AIS of 50% of all neurons. When present at the AIS, membrane GFP signal only occupied 0.16% of AIS surface (∼30-times lower than at dendritic membrane sites; 4.8% occupancy, **Fig. 4A,B**). A subsequent total GFP staining revealed a diffuse pattern in the neuron, including in the AIS, as well as bright and dense puncta in the postsynaptic density (**fig. S6A**).

**Fig. 4.**
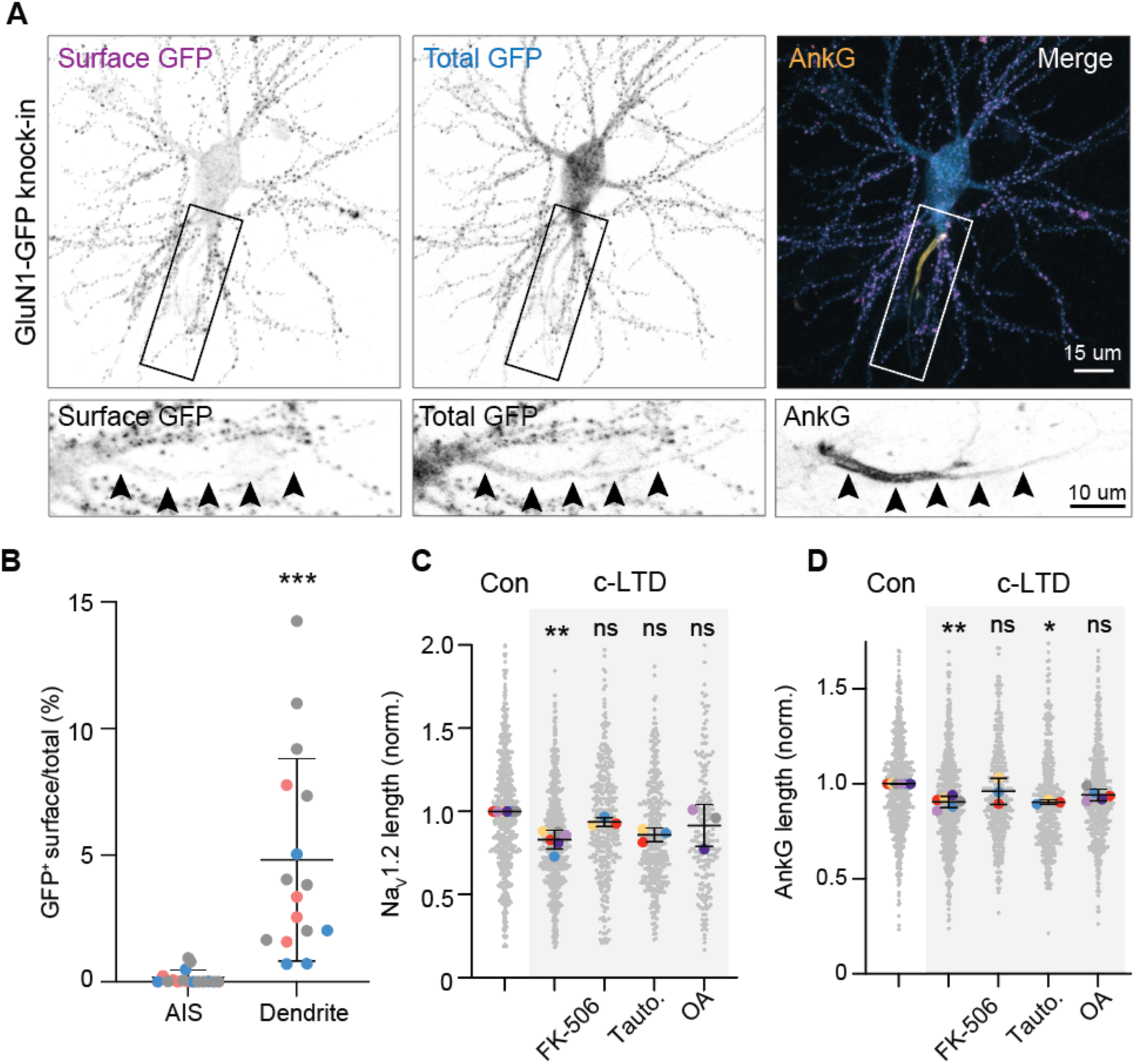
AIS plasticity is triggered by the activation of synaptic NMDARs and relies on calcineurin (A) GluN1-GFP knock-in neurons stained for extracellular GFP, total GFP and AnkG. Scale bars are 15 µm and 10 µm in the zooms. Arrows show the trajectory of the AIS and axon. **(B)** Percentage of the AIS or dendrites surface positive for extracellular GFP. Wilcoxon’s test, ***p = 0.001, *n* = 16 neurons from 3 independent cultures. (**C,D**) Na_V_1.2 and AnkG relative lengths in c-LTD and in the presence of FK-506, Tautomycetin (Tauto) or Okadaic Acid (OA). *N* = 3 to 6 cultures, Kruskal-Wallis test. For Na_V_1.2, c-LTD **p = 0.0024, FK-506 ns p = 0.93, Tauto ns p = 0.06, OA ns p = 0.82. For AnkG: c-LTD **p = 0.004, FK-506 ns p = 0.99, Tauto *p = 0.015, OA ns p = 0.28.

The low abundancy of NMDARs at the AIS suggests that the effect of NMDA application might be mediated via NMDARs at somato-dendritic and synaptic sites. To test this idea experimentally, we used MK-801, blocking glutamate-bound NMDARs (E. H. Wong et al., 1986; Woodruff et al., 1987), often used to distinguish extrasynaptic from synaptic NMDARs. MK-801 applied 5 min before and during NMDA application blocked the shortening of both AnkG and Na_V_1.2, indicating that activation of synaptic NMDARs is required to trigger AIS plasticity (**Fig. S6B,C**). Taken together, the absence of NMDARs inserted at the AIS membrane and the requirement of activated NMDARs for the induction of AIS shortening reveals that the activation of synaptic NMDAR is the trigger for rapid AIS plasticity.

Which signaling pathway links synaptic NMDAR activation with AIS plasticity? We observed that NMDA application leads to LTD in acute slices (**Fig. 1C**) but also induces a strong, membrane depolarization and action potential generation lasting for several minutes (**Fig. 1D**). To test whether induction of synaptic LTD alone suffices to drive AIS plasticity we evoked LTD electrically (e-LTD, SC stimulation, 900 pulses @1 Hz, **fig. S6D,E**). Although we observed strong LTD (EPSC peak amplitude reduction ∼48% e-LTD vs 8% control) we did not observe AIS length changes, neither in control nor in e-LTD (**fig S6F,G**). We next hypothesized that the postsynaptic depolarization and associated action potentials after NMDA application sufficed to trigger AIS plasticity. To test this conjecture directly, we reproduced the NMDA-induced depolarization by injecting positive DC current for 5 min, keeping the neuron above threshold (∼ –40 mV) and generating action potential firing. However, the results showed that the AIS length remained constant (**fig. S6H**). Altogether these results suggest that the combination of synaptic depression as well as the postsynaptic firing is required to trigger NMDAR-mediated Na_V_1.2 removal from the distal AIS.

We then sought to identify the molecular signaling cascade mediating rapid AIS remodeling. The induction of c-LTD is known to activate numerous phosphatases and interestingly some of these have also been shown to be implicated in AIS plasticity, like calcineurin (Evans et al., 2013). To test the involvement of calcineurin and other phosphatases, we applied NMDA in the presence of diverse pharmacological inhibitors to assess their ability to prevent the shortening of Na_V_1.2 and AnkG (**Fig. 4C,D**, respectively). Inhibiting calcineurin (FK-506, 50 mM) or PP2A (with low dose of Okadaic Acid, OA, 2 nM) blocked the shortening of both AnkG and Na_V_1.2 induced by NMDAR activation. Inhibition of PP1 (Tautomycetin, 10 µM) prevented the shortening of Na_V_1.2 but not of AnkG (**Fig. 4C,D**). Thus, these results show that dephosphorylation is an important process in the removal of Na_V_1.2 at the AIS and reveal a common signaling pathways leading to the expression of LTD at synapses and the AIS after activation of NMDAR.

### AIS shortening is mediated by clathrin-mediated endocytosis of Nav1.2 from the distal AIS

How are Na_V_ channels removed from the distal AIS during NMDAR-mediated AIS plasticity? We pharmacologically inhibited common signaling pathways known to promote protein removal and/or degradation (**Fig. 5A,B**, and **fig.S7**). We first tested the role of calpains, a family of calcium-dependent proteases. Although calpains have been shown to be involved in stress-mediated AIS disassembly (Benned-Jensen et al., 2016; Benusa, George, Sword, DeVries, & Dupree, 2017; Clark, Sword, & Dupree, 2017; Del Puerto et al., 2015; Schafer et al., 2009) inhibition with MDL 28170 during induction of c-LTD neither blocked the ∼15% shortening of Na_V_1.2 nor AnkG (**Fig. 5A,B**). Since it was shown previously that AIS proteostasis can be modulated by local Ecm29-targeted proteasome (M. Lee et al., 2020) we then tested the role of proteasome-mediated degradation. Interestingly, induction of c-LTD in the presence of the proteasome-blocker MG-132 prevented the shortening of AnkG, but not Na_V_1.2 (**Fig. 5B,C** and **fig. S7A**).

**Fig. 5.**
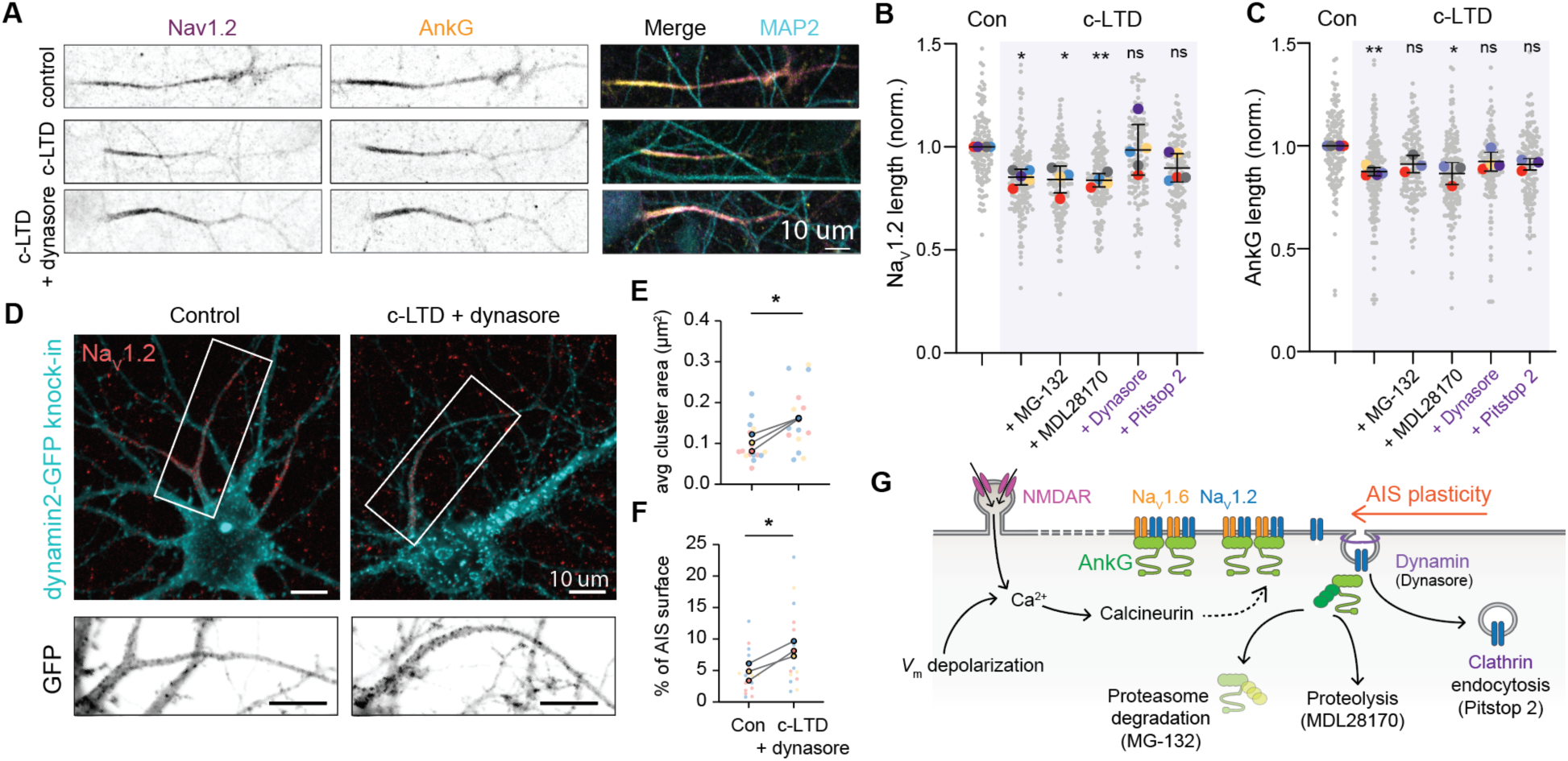
Plasticity-induced Na_V_1.2 removal is mediated by endocytosis. (**A**) Immunostaining for Na_V_1.2, AnkG and MAP2 of DIV14 neurons in control conditions and c-LTD with or without the dynamin inhibitor dynasore. (**B**) Na_V_1.2 and (**C**) AnkG relative length following c-LTD and in the presence of MG-132, MDL2870, dynasore and Pitstop 2, *N* = 3-6 independent experiments and *n* >100 neurons per condition per experiment. Kruskal-Wallis test; c-LTD: Nav1.2 *p = 0.019, AnkG **p = 0.001, c-LTD + MG-132: Nav1.2 *p = 0.036, AnkG ns p = 0.21, c-LTD + MDL28170: Na_V_1.2 **p = 0.005, AnkG *p = 0.018, c-LTD + dynasore: Na_V_1.2 ns p > 0.99, AnkG ns p = 0.38, c-LTD + Pitstop 2: Na_V_1.2 ns p = 0.068, AnkG ns p = 0.39. (**D**) Immunostaining for dynamin2 reveals increased NMDA-induced dynamin clusters in the AIS. Scale bar is 20 µm. (**E, F**) Population analysis of dynamin2 cluster areas and dynamin surface in the Na_V_1.2-positive AIS. In (E): paired t-test, *p = 0.035 and in (F): paired t-test, *p = 0.033. N = 3 independent experiments, from 4 to 7 neurons per condition per experiment. (**G**) Summary scheme and working model. Synaptic NMDARs activation and APs generate local Ca^2+^ influx activating downstream effectors, including calcineurin and lead to AnkG degradation by the UPS and local recruitment of dynamin2 responsible for Na_V_1.2 channels internalization.

Finally, we addressed the involvement of endocytosis in controlling Na_V_1.2 re-distribution during plasticity. Na_V_1.2 and Na_V_1.6 channels have been reported to contain endocytic motifs which are important for their stabilization at the AIS membrane (Fache et al., 2004; Sole et al., 2019). Consistent with these observations, the application of two distinct blockers for endocytosis, either by inhibiting dynamin GTPase activity (dynasore, 80 µM) or clathrin (Pitstop 2, 20 µM), abolished the NMDA-induced shortening of Na_V_1.2 and AnkG (**Fig. 5C, Fig. S7A, B**), showing that endocytosis is required for AIS plasticity. Furthermore, application of dynasore alone did not affect AnkG length (**Fig. S7C**).

These blockers also inhibit the expression of c-LTD at the PSD (Beattie et al., 2000; Carroll et al., 1999), thus to address the involvement of endocytosis specifically at the AIS during plasticity, we first assessed the presence at the AIS of two members of the endocytic machinery, dynamin2 (**Fig. 5D**) and clathrin light chain (**fig. S7D**) using CRISPR/Cas9 GFP knock-ins (Willems et al., 2020). We detected GFP puncta throughout the transfected neurons including at the AIS, stained with Na_V_1.2. We then applied dynasore during c-LTD to assess whether a pool of endocytic structures would be specifically recruited to and retained at the AIS during c-LTD (Leyton-Puig et al., 2017). We observed a significant increase of the relative surface area covered by dynamin2-GFP at the AIS (∼1.5 fold), and of the average cluster area after c-LTD (∼1.5 fold) in the presence of dynasore (**Fig. 5D-F**). No change was detected in the pattern of clathrin light chain (fig. S5). These results suggest a model in which dynamin2 is rapidly recruited to the AIS during NMDAR-mediated plasticity. Altogether, our data show that activity-dependent Na_V_1.2 channel removal at the distal AIS is mediated by clathrin-mediated endocytosis.

## Discussion

Using real-time imaging of novel endogenous labels for the scaffold protein AnkG and Na_V_ channels, we find that the chemical induction of synaptic c-LTD, characterized by increased neuronal depolarization and a rapid depression of synaptic responses, triggers structural AIS plasticity within minutes via local internalization of Na_V_1.2 channels and in a dynamin-dependent manner (Fig. 5). The rapid disassembly of the distal AIS shares multiple molecular mechanisms with the expression of synaptic LTD, from the role of clathrin-dependent endocytosis in the removal of AMPARs from the PSD (Beattie et al., 2000; Man et al., 2000), to the requirement of calcineurin-mediated dephosphorylation (H. K. Lee, Takamiya, He, Song, & Huganir, 2010; Oh, Derkach, Guire, & Soderling, 2006). It would be interesting to investigate whether calcineurin directly dephosphorylates Na_V_1.2, and address how this affects Na_V_1.2 binding to AnkG and membrane dynamics of the channel. Na_V_1.2 bears an endocytic motif in the II-III loop, which is masked by the interaction with AnkG, and thereby could block the internalization of the channel at the AIS (Fache et al., 2004). Interaction with AnkG alone however, might not be sufficient to prevent internalization, as shown for the MAP1B-binding mutant Na_V_1.6 channel, which is still capable to bind to AnkG, but accumulates less at the AIS membrane because it undergoes endocytosis (Sole et al., 2019). The third mechanism shared with synaptic LTD is the contribution of the ubiquitin proteasome degradation system (UPS), which controls PSD-95 and AKAP150 levels during LTD (Cheng et al., 2020; Colledge et al., 2003). Similarly, we observed that that blocking the UPS during LTD prevents the shortening of AnkG (Fig. 5B), indicating that AIS remodeling could also rely on protein degradation. A role for the UPS in the AIS proteostasis, as well as for 190-kDa AnkG stability at the spine has previously been reported (M. Lee et al., 2020; Yoon et al., 2020). If AnkG is targeted by the UPS, this is not the case for Na_V_1.2 (Fig. 5A,B) and in future studies it would be important to assess the fate of endocytosed Na_V_1.2 channels. Endocytosed Na_V_1.2 channels could be targeted to lysosomal degradation or may undergo local recycling, which could provide a ‘ready-releasable’ recycling pool of Na_V_ channels, and it is tempting to speculate that such pool could be used during rapid AIS remodeling. Once endocytosed, the targeting of Na_V_1.2 for recycling *versus* degradation might rely on the duration and concentration of cytoplasmic Ca^2+^. In the case of excitotoxic events, the AIS is disrupted in an irreversible manner. Indeed, applying high concentration of glutamate causes the endocytosis and subsequent degradation of AIS K_V_7.2/7.3 channels, in an NMDAR-dependent manner (Benned-Jensen et al., 2016).

Our results suggests that for rapid AIS shortening the temporal coincidence of synaptic activation and postsynaptic depolarization is a requirement: AnkG-GFP remained stable with low-frequency presynaptic stimulation inducing e-LTD as well as during postsynaptic depolarization and associated AP firing (Fig. S3F-H). These data are compatible with a model in which temporally correlated activation of presynaptic inputs with postsynaptic depolarization, is required to trigger rapid AIS plasticity. While these conditions are experimentally achieved with exogenous NMDA application (depolarizing spines and dendritic branches as well as the axonal membrane via minutes-long AP generation (this study and (H. K. Lee et al., 1998)), how spike-timing dependent plasticity or other physiologically-relevant stimuli induce AIS plasticity and ion channel endocytosis/insertion will require further studies. Rapid AIS plasticity mediated by correlated synaptic activation and action potential generation may provide a mechanistic explanation for the NMDAR-dependent synergistic scaling of the AP threshold and synaptic responses during LTP (Xu, Kang, Jiang, Nedergaard, & Kang, 2005). The AnkG-GFP transgenic mouse and Na_V_ live markers presented here enables experiments to correlate cytoplasmic Ca^2+^ imaging with ion channel membrane dynamics and establish how spatiotemporal patterns of activity-dependent intracellular Ca^2+^ levels change links with AIS Na_V_1.2 endocytosis. Rapid spike threshold changes are a hallmark of intrinsic plasticity and observed across many species, learning paradigms and intrinsic plasticity protocols (Zhang & Linden, 2003). Since activity-dependent changes of the AIS provides a critical hub for intrinsic plasticity, cooperation with synaptic plasticity via common molecular pathways may represent a powerful mechanism to change the neuronal ensemble activity and engram formation during learning and memory.

## Materials and Methods

### Animals & Ethics

Double-floxed AnkG-GFP mice (B6;129SV-ank3tm1DCI/HD) were generated by Vann Bennett (previously: Duke University, North Carolina, USA) and Paul Jenkins (University of Michigan, Michigan, USA) and a kind gift by Maren Engelhardt (Johannes-Kepler-University, Linz, Austria). Mice of both sexes were kept at a 12h light-dark cycle (lights on at 07:00, lights off at 19:00) with ad libitum food and water. All animal experiments were performed in compliance with the European Communities Council Directive 2010/63/EU effective from 1 January 2013 and with Dutch national law (Wet op de Dierproeven, 1996). The experimental design and ethics were evaluated and approved by the national committee of animal experiments (CCD) applications number AVD 80100 2017 2426 (Netherlands Royal Academy of Arts and Sciences, KNAW) and AVD 10800 2017 3404 (Utrecht University). The animal experimental protocols were designed to minimize suffering and approved and monitored by the KNAW animal welfare body (NIN 14.49, NIN 12.13, NIN 20.21.07.

### AAV injections

Between 4 to 6 weeks of age mice were anesthetized with isoflurane and head-fixed in a stereotactic apparatus (Kopf instruments). Body temperature was kept at 35 °C with a heating pad. A glass pipette was inserted bilaterally through a small craniotomy (stereotactic coordinates: 3.0 mm P, 3.0 mm L, 2.0-1.5 mm depth). 50 nl of undiluted pENN.AAV.CamKII 0.4.Cre.SV40 (Plasmid #105558, AAV5, Addgene) were injected into the CA1 area of the hippocampus. Cre-recombinase flips the orientation of a cassette containing both the *Ank3* last exon and the coding sequence of eGFP, thus inducing expression of Cre-dependent AnkG-GFP fusion protein. Afterwards the skin was closed with sutures and mice received 1 mg/kg BW meloxicam (Metacam®, Boehringer-Ingelheim, Germany). s.c. as post-operative pain medication. Mice were allowed to recover for at least 2 weeks prior to sacrifice, to allow for sufficient virus expression.

### Acute slice preparation

Between 6 to 8 weeks of age, mice were sacrificed for preparation of acute hippocampal brain slices. Mice were deeply anesthetized by application of pentobarbital s.p. (60 mg/kg BW). They were perfused with ice-cold oxygenated ACSF (125 mM NaCl, 25mM NaHCO_3_, 1.25 mM NaH_2_PO_4_, 3 mM KCl, 25 mM Glucose, 1 mM CaCl_2_, 6 mM MgCl_2_, and 1 mM kynurenic acid, saturated with 95% O_2_ and 5% CO_2_, pH 7.4) and subsequently decapitated. The brain was swiftly removed and submerged in ice-cold ACSF (composition see above). 300 μm thick transverse acute hippocampal slices were cut using a Leica VT 1200S vibratome (Leica Biosystems, Wetzlar Germany). Slices were allowed to recover for at least 35 min (60 min for e-LTD experiments) at 35 °C and subsequently kept in a holding chamber at room temperature until experiments commenced. Recordings and incubation experiments were carried out at ∼32 °C.

### Acute slice experiments

After sectioning and recovery, the slices were gently transferred onto cell culture inserts (Millicell, 30 mm, Merck) in a small petri dish that was constantly perfused with ACSF (125 mM NaCl, 25 mM NaHCO_3_, 1.25 mM NaH_2_PO_4_, 3 mM KCl, 25 mM Glucose, 2 mM CaCl_2_, 1.3 mM MgCl_2_ saturated with 95% O_2_ and 5% CO_2_, pH 7.4) at 6 ml/min. After transfer, the chamber was perfused with 20 μM NMDA (Sigma-Aldrich) in ACSF for 3 minutes, directly followed by perfusion with 100 μM APV (Alfa Aesar) for 3 minutes to prevent prolonged NMDA receptor opening. For control slices, treatment solely consisted of perfusion with 100 μM APV for 3 minutes to exclude potential effects of APV on AIS length. Subsequently, slices were fixated using 4% paraformaldehyde (PFA, in 0.1 M PBS) for 20 minutes at two time-points: before perfusion, as well as after 30 minutes post treatment. NMDA-treated slices and control slices were taken from corresponding hemispheres at matching CA1 locations (dorsoventral axis) to exclude potential effects of location on AIS length. Slices were then subjected to multichannel immunofluorescent staining. Slices were incubated in blocking buffer (10% NGS, 1% Triton in 0.1 M PBS) for a minimum of 90 minutes at room temperature, followed by incubation in primary antibodies overnight (5% NGS, 1% Triton in 0.1 M PBS). Primary and secondary antibodies and concentrations were the same as used for cryosections (see above). Following washing, the slices were incubated with secondary antibodies for a minimum of 4 hours in the dark. Streptavidin Alexa-633 conjugate (1:1000; Invitrogen) was added for the detection of biocytin filled neurons. Subsequently, slices were washed and mounted using fluorescence-preserving mounting medium (Vectashield, Vector laboratories). Confocal images of the AIS were acquired using a Leica SP8 confocal microscope (Leica Microsystems, Wetzlar Germany). A 40× objective (oil immersion, NA 1.4) was used for population analysis of AIS length. To ensure that the entire AIS structure was in focus, multiple images in the z-dimension were projected to one image (Maximum intensity projection). Each image section was spaced at 0.5 μm at a total depth of up to 25 μm. Images were taken at 1024 × 1024 resolution, resulting in resolution of 0.27 μm/ pixel. Individual AIS length was determined using the morphometrical software analysis tool AISuite (https://github.com/jhnnsrs/aisuite2). Segments were individually traced, and the threshold determining start and end of each individual segment was set to be 35% of the maximum fluorescent signal of a single AIS.

### Electrophysiology

Slices were transferred to an upright microscope (BX61WI, Olympus Nederland BV) and constantly perfused with oxygenated ACSF: 125 mM NaCl, 25 mM NaHCO_3_, 1.25 mM NaH_2_PO_4_, 3 mM KCl, 25 mM Glucose. Depending on the recording, 2 mM CaCl_2_, 1.3 mM MgCl_2_ and 1 mM sodium ascorbate were added (c-LTD) or 2.5 mM CaCl_2_ and 1 mM MgCl_2_ (e-LTD and depolarisation experiments) were added. The chamber was perfused at a rate of 3 mL/min. Neurons were visualized with a 40× water immersion objective (Achroplan, NA 0.8, IR 40×/0.80 W, Carl Zeiss Microscopy) with infrared optics and oblique contrast illumination. Patch-clamp recordings were performed from CA1 pyramidal neurons with a GFP positive AIS identified during a search in epifluorescence mode and visualizing GFP with the 470 nm laser line of a laser diode illuminator (LDI-7, 89 North, USA). Patch pipettes were pulled from borosilicate glass (Harvard Apparatus, Edenbridge, Kent, UK) pulled to an open tip resistance of 4–5 MΩ and filled with intracellular solution containing: 130 mM K-Gluconate, 10 mM KCl, 10 mM HEPES, 4 mM Mg-ATP, 0.3 mM Na2-GTP, and 10 mM Na2-phosphocreatine (pH 7.25, ∼280 mosmol). Atto-594 (20 µM, Atto-tec, Germany) and biocytin (3 mg/ml, Sigma-Aldrich) were routinely added to the intracellular solution to allow for live- and post-hoc confirmation of colocalization of the AIS with the recorded neuron.

Recordings were performed with a Axopatch 200B (Molecular Devices). Signals were analogue low-pass filtered at 10 kHz (Bessel) and digitally sampled at 50 kHz using an A-D converter (ITC-18, HEKA Elektronik Dr. Schulze GmbH, Germany) and the data acquisition software Axograph X (v.1.5.4, Axograph Scientific, NSW, Australia). Bridge-balance and capacitances were fully compensated in current clamp. Series resistance was compensated to > 75% for voltage-clamp recordings. Once a stable recording was established, action potential threshold was recorded at the start (and end) of recordings by injecting 3 ms pulses at increasing current steps (10 pA) until an action potential was generated. For LTD recordings, an extracellular glass bipolar stimulation electrode was positioned above the Schaffer collaterals, to stimulate presynaptic inputs to CA1. The external electrode was tuned to evoke half of the maximum currents in the recorded cell. Initially, > 5 min. of baseline was recorded, and all postsynaptic currents were normalized to the mean evoked current amplitude during baseline. For chemical LTD (c-LTD) experiments, LTD was evoked by 3 min. application of 20 μM NMDA (Sigma-Aldrich), based on the protocols by (Kamal, Ramakers, Urban, De Graan, & Gispen, 1999; H. K. Lee et al., 1998). In initial experiments we observed excessive depolarization with NMDA and therefore applied subsequently 100 μM APV (Alfa Aesar) to rapidly antagonize NMDAR-mediated currents. Thus, APV limited and temporally controlled the duration of the glutamatergic receptor activation. Neuronal survival was further improved by adding sodium ascorbate and increase MgCl_2_ to 1.3 mM.

The AIS was imaged using an Olympus FV1000 confocal microscope (Olympus corporation, Tokyo, Japan) controlled by the imaging software FV10-ASW (Ver.03.00). Z-stacks were acquired at a speed of 12.5 µs/ pixel and a resolution of 800 × 800 pixels in 1 µm steps at 1– 3× zoom. To reduce phototoxicity and minimize bleaching effects of the laser, z-stacks of AIS were taken at four time points: i) before baseline, ii) after baseline/ before treatment, iii) 30 min. post treatment, iv) 60 min. post treatment. AIS length was determined with the “Measure ROIs” plugin (https://github.com/cleterrier/Measure_ROIs) in Fiji (ImageJ). Fluorescence intensity thresholds for the detection of AIS beginning and end were adjusted between 10% and 35% depending on the signal-to-noise ratio, but kept consistent for an individual AIS time series. For some c-LTD (*n* = 12/19) and APV control (*n* = 8/17) experiments, cells were only imaged but not patched. Several cells per slice were imaged simultaneously to increase data yield. In these cases, healthy neurons were picked based on visual criteria (low contrast edges, no swelling or shrinking).

### Primary neuronal cultures and transfection

Primary hippocampal neurons cultures were prepared from embryonic day 18 rat brains (both genders). Cells were plated on coverslips coated with poly-L-lysine (37.5 μg/mL, Sigma-Aldrich) and laminin (1.25 μg/mL, Roche Diagnostics) at a density of 100,000/well of 12-wells plates. Neurons were first cultured in Neurobasal medium (NB) supplemented with 2% B27 (GIBCO), 0.5 mM glutamine (GIBCO), 15.6 μM glutamate (Sigma-Aldrich), and 1% penicillin/streptomycin (GIBCO) at 37 °C in 5% CO_2_. From DIV1 onwards, half of the medium was refreshed weekly with BrainPhys medium (BP) supplemented with 2% NeuroCult SM1 neuronal supplement (STEMCELL Technologies) and 1% penicillin/streptomycin.

Hippocampal neurons were transfected using Lipofectamine 2000 (Invitrogen). Briefly, DNA (1.8 μg/well, of a 12-wells plate) was mixed with 3.3 μL of Lipofectamine 2000 in 200 μL NB, incubated for 20 min, and then added to the neurons in Neurobasal at 37°C in 5% CO_2_ for 1 hr. Then, neurons were washed with Neurobasal and transferred back to their original medium. Transfection of knock-in constructs was performed at DIV3.

### DNA Constructs

The knock-in constructs of GluN1-GFP and clathrin light chain A-GFP (Willems et al., 2020), and dynamin2-GFP (Catsburg, Westra, van Schaik, & MacGillavry, 2022) were previously described. The knock-in construct for Nav1.2 was cloned in the pORANGE (Willems et al., 2020) and the gRNA used was: GGACAAAGGGAAAGATATCA. The GFP tag was added in the C-Terminal region (4aa before the STOP codon), mimicking the tagging approach used for Na_V_1.6 (Akin et al., 2015; Gasser et al., 2012) or Nav1.2 (Liu et al., 2022), which was shown to not alter channel function nor localization.

### Antibodies

The following antibodies were used in this study: Na_V_1.2 (NeuroMab/Antibodies Incorporated #75-024), Na_V_1.6 (Alomone Labs #ASC-009 and NeuroMab/Antibodies Incorporated #73-026), Pan-Nav (Sigma, #S8809), AnkG (Life technologies #33-8800, Neuromab/Antibodies Incorporated #75-146 and Synaptic systems #386-005), βIV-spectrin (Biotrend, provided by Maren Engelhardt, Johnnes-Kepler-University, Linz, Austria), MAP2 (Abcam/Bio Connect #ab5392), GFP (MBL International/Sanbio #598 and Abcam #ab13970).Corresponding secondary antibodies Alexa-conjugated 405, 488, 568, 594 or 647 goat anti-mouse, anti-mouse IgG1/IgG2a, anti-rabbit, anti guinea-pig or anti-chicken were used (Life Technologies), as well as streptavidin Alexa-633 conjugate (Life Technologies) for the detection of biocytin filled neurons.

### Pharmacological treatments and c-LTD experiment

MG-132 (Calbiochem, #474790), MDL 28170 (Tocris Bioscience, #1146/10), MK-801 (Tocris Bioscience, #924), Okadaic Acid (Tocris Bioscience, #1136), Tautomycetin (Tocris Bioscience, #2305), FK-506 (Tocris Bioscience, #3631), Dynasore (Tocris Bioscience, #2897), PitStop2 (Abcam, #ab120687), APV (Tocris Bioscience, #0106), NMDA (Sigma #3263). All chemicals were diluted in a modified Tyrode’s solution (pH 7.4) containing 25 mM HEPES, 119 mM NaCl, 2.4 mM KCl, 2 mM CaCl2, 2 mM MgCl2, 30 mM glucose. For c-LTD experiments, neurons were first incubated for 5 min in modified Tyrode’s solution, then NMDA 2X (diluted in modified Tyrode’s at 100 μM) was added to the wells, alternatively, Tyrode’s solution was applied as a control. After 4 min, neurons were dipped in Tyrode’s solution to wash them and returned to their original medium. c-LTD combined with live imaging was performed the same way, all incubation and washing steps were performed on stage. Triton extraction experiment: Live DIV14 hippocampal neurons were incubated for 5 min at RT in 0.25 % Triton X-100 preheated at 37 °C. Neurons were fixed directly after extraction.

### Immunocytochemistry

For immunocytochemistry of cultured neurons, cells were fixed for 6 min with warm paraformaldehyde (4%)-sucrose (4%). Primary antibodies were incubated overnight at 4 °C in GDB buffer (0.2% BSA, 0.8 M NaCl, 0.5% Triton X-100, 30 mM phosphate buffer, pH 7.4). After 3 washes in PBS, secondary antibodies were incubated in the same buffer for 1hr at RT. Extracellular staining of GluN1 was performed using a first incubation step with the anti-GFP antibody O/N at 4 °C in GDB without Triton, followed by 3 washes in PBS and incubation with an Alexa568 secondary antibody in GDB without Triton for 2 h at RT. Then anti-GFP and anti-AnkyrinG antibodies were incubated in GDB buffer containing Triton O/N at 4 °C, washed 3 times and corresponding secondary antibodies conjugated with Alexa405 and 647 were incubated in the same buffer for 1hr at RT. Coverslips were mounted using Vectashield (Vectorlabs).

### Immunohistochemistry

To examine the expression of AnkG-GFP mice were deeply anesthetized by application of pentobarbital s.p. (60 mg/kg BW) prior to transcardial perfusion with PBS for 5 minutes followed by 4% paraformaldehyde (PFA, in 0.1M PBS) for 5 minutes. Brains were postfixed in 4% PFA for 1 hour at room temperature (RT). After three sucrose cryoprotection steps (1 h in 10% RT, overnight in 20%, 4°C, overnight in 30% sucrose, 4°C) brains were trimmed to sectioning block in coronal orientation and frozen in embedding medium (Tissue-Tek®, Sakura) until further processing. Brains were cut with a cryostat (CM3050 S, Leica) to sections of 40 µm thickness. Slices were incubated in blocking buffer (10% normal goat serum (NGS), 0.5% Triton in 0.1 M PBS) for a minimum of 90 minutes at RT, followed by incubation in primary antibodies overnight (5% NGS, 0.5% Triton in 0.1 M PBS). Following washing steps, the slices were incubated with secondary antibodies for a minimum of 90 minutes in the dark. Subsequently, slices were washed and mounted using fluorescence-preserving mounting medium (Vectashield, Vector laboratories).

### Image acquisition

Primary neurons were imaged using a LSM700 confocal laser-scanning microscope (Zeiss) with a Plan-Apochromat 63× NA 1.40 oil DIC, EC Plan-Neofluar 40× NA1.30 Oil DIC and a Plan-Apochromat 20× NA 0.8 objective. Z-stacks were acquired with 0.5 μμm steps and maximum projections were done from the resulting stack. Acquisition settings were kept the same for all conditions within an experiment.

### gSTED microscopy

Gated STED (gSTED) imaging was performed with a Leica TCS SP8 STED 3X microscope using a HC PL APO 100 × / 1.4 oil immersion STED WHITE objective. The 594 and 647 nm wavelengths of pulsed white laser (80 MHz) were used to excite the Alexa594 and the Alexa647 secondary antibodies. Both Alexa594 and Alexa647 were depleted with the 775 nm pulsed depletion laser. Fluorescence emission was detected using a Leica HyD hybrid detector with a time gate of 0.3 ≤ tg ≤ 6 ns.

### Live cell imaging and FRAP experiments

Live-cell imaging experiments were performed in an inverted microscope Nikon Eclipse Ti-E (Nikon), equipped with a Plan Apo VC 100x NA 1.40 oil, a Plan Apo VC 60× NA 1.40 oil and a Plan Apo VC 40× NA 1.30 oil objectives (Nikon), a Yokogawa CSU-X1-A1 spinning disk confocal unit (Roper Scientific), a Photometrics Evolve 512 EMCCD camera (Roper Scientific) and an incubation chamber (Tokai Hit) mounted on a motorized XYZ stage (Applied Scientific Instrumentation) which were all controlled using MetaMorph (Molecular Devices) software. Coverslips were mounted in metal rings and imaged using an incubation chamber that maintains temperature and CO_2_ optimal for the cells (37 °C and 5% CO_2_). Live imaging was performed in full conditioned medium. Time-lapse live-cell imaging of Na_V_1.2 knock-in neurons was performed with a frame rate of 1 acquisition per minute for the FRAP experiments and 1 acquisition every 7.5 minutes for the c-LTD experiments. FRAP experiments were carried out with an ILas FRAP unit (Roper Scientific France/ PICT-IBiSA, Institut Curie). Local photobleaching of Na_v_1.2-GFP in the proximal and distal AIS was performed using ROIs of 30 × 30 pixels (∼ 25 μm^2^) and recovery was monitored for 45 min, with an image acquisition rate of 1 image per minute.

### Image processing, quantifications, and statistical analysis

Movies and images were processed using Fiji (https://imagej.net/software/fiji/) (Schindelin et al., 2012). Kymographs were generated using the ImageJ plugin KymoResliceWide v.0.5 https://github.com/ekatrukha/KymoResliceWide. The AIS position (start and end points), as well as fluorescence intensity of AIS proteins were measured using Christophe Leterrier’s plugin Pro_Feat_Fit (https://github.com/cleterrier/Measure_ROIs), and with the following parameters: Small detection, starting point at 50% and end-point at 35% of the max fluorescence intensity. The autocorrelation coefficients were obtained using Christophe Leterrier’s plugin ‘Autocorrelation’ (https://github.com/cleterrier/Process_Profiles). GluN1, Clathrin light chain and Dynamin2 spots quantifications were performed using the Spot Detector plug-in built-in in Icy software (https://icy.bioimageanalysis.org) (de Chaumont et al., 2012). Quantifications parameters were adjusted per experiment but kept similar for all conditions within each experiment. FRAP recovery was quantified as the percentage of fluorescence recovery depending on the initial fluorescence intensity and taking into account the acquisition photobleaching. A bleaching control was selected outside of the bleached areas and fluorescent signals in the ROIs was set to 100% based on the mean intensity of the first 4 frames (4 min) before bleaching in the same region and was set to 0% directly after as described previously (Phair, Gorski, & Misteli, 2004). The average recovery rate was calculated by averaging the values of the last 5 points of fluorescence intensity.

All statistical details of experiments, including the exact values of n, and statistical tests performed, are shown in Figures and Figure Legends. n represents the number of neurons analyzed, and N the number of independent experiments. All statistical analyses were performed using Prism9 (GraphPad Software).

Significance was defined as: ns-not significant, *p < 0.05, **p < 0.01, and ***p < 0.001. Normality of the data was determined by a D’Agostino and Pearson’s test and parametric two-tailed paired or unpaired t tests or non-parametric Mann Whitney or Wilcoxon tests were applied when comparing two groups. For more than two groups, one- or two-ways ANOVA were used followed by a Tukey’s multiple comparison test. When values were missing (i.e. during live imaging, when the 60 minute time point could not be imaged due to loss of the recorded cell), a mixed-effects analysis followed by Šídák’s multiple comparisons was applied. For all experiments in cultured neurons, the statistical significance was tested on the average values of each independent experiment, except for quantification of knock-in neurons. At least 3 independent experiments were performed. For normalized data, each value was normalized to the average value of the control condition within the same experiment.

## Acknowledgments

The authors are indebted to the support by the neuron culture team and the imaging facility at Utrecht University. We would like to thank Dr Harold MacGillavry for the great discussions and help throughout the project.

## Author contributions

Conceptualization: AF

Methodology: AF, MHPK

Investigation: AF, NJ, JTB, JJ, TL, NP

Visualization: AF, NJ, MHPK

Funding acquisition: AF, MHPK

Project administration: AF, MHPK

Supervision: AF, MHPK

Writing – original draft: AF, NJ, MHPK

Writing – review & editing: AF, NJ, MHPK, CCH

## References

Akin, E. J., Sole, L., Dib-Hajj, S. D., Waxman, S. G., & Tamkun, M. M. (2015). Preferential targeting of Nav1.6 voltage-gated Na+ Channels to the axon initial segment during development. PLoS One, 10(4), e0124397. doi:10.1371/journal.pone.0124397

Beattie, E. C., Carroll, R. C., Yu, X., Morishita, W., Yasuda, H., von Zastrow, M., & Malenka, R. C. (2000). Regulation of AMPA receptor endocytosis by a signaling mechanism shared with LTD. Nat Neurosci, 3(12), 1291–1300. doi:10.1038/81823

Benned-Jensen, T., Christensen, R. K., Denti, F., Perrier, J. F., Rasmussen, H. B., & Olesen, S. P. (2016). Live Imaging of Kv7.2/7.3 Cell Surface Dynamics at the Axon Initial Segment: High Steady-State Stability and Calpain-Dependent Excitotoxic Downregulation Revealed. J Neurosci, 36(7), 2261–2266. doi:10.1523/JNEUROSCI.2631-15.2016

Benusa, S. D., George, N. M., Sword, B. A., DeVries, G. H., & Dupree, J. L. (2017). Acute neuroinflammation induces AIS structural plasticity in a NOX2-dependent manner. J Neuroinflammation, 14(1), 116. doi:10.1186/s12974-017-0889-3

Berger, S. L., Leo-Macias, A., Yuen, S., Khatri, L., Pfennig, S., Zhang, Y., … Salzer, J. L. (2018). Localized Myosin II Activity Regulates Assembly and Plasticity of the Axon Initial Segment. Neuron, 97(3), 555–570 e556. doi:10.1016/j.neuron.2017.12.039

Carroll, R. C., Beattie, E. C., Xia, H., Luscher, C., Altschuler, Y., Nicoll, R. A., … von Zastrow, M. (1999). Dynamin-dependent endocytosis of ionotropic glutamate receptors. Proc Natl Acad Sci U S A, 96(24), 14112–14117. doi:10.1073/pnas.96.24.14112

Catsburg, L. A., Westra, M., van Schaik, A. M., & MacGillavry, H. D. (2022). Dynamics and nanoscale organization of the postsynaptic endocytic zone at excitatory synapses. Elife, 11. doi:10.7554/eLife.74387

Chand, A. N., Galliano, E., Chesters, R. A., & Grubb, M. S. (2015). A distinct subtype of dopaminergic interneuron displays inverted structural plasticity at the axon initial segment. J Neurosci, 35(4), 1573–1590. doi:10.1523/JNEUROSCI.3515-14.2015

Cheng, W., Siedlecki-Wullich, D., Catala-Solsona, J., Fabregas, C., Fado, R., Casals, N., … Minano-Molina, A. J. (2020). Proteasomal-Mediated Degradation of AKAP150 Accompanies AMPAR Endocytosis during cLTD. eNeuro, 7(2). doi:10.1523/ENEURO.0218-19.2020

Christie, J. M., & Jahr, C. E. (2009). Selective expression of ligand-gated ion channels in L5 pyramidal cell axons. J Neurosci, 29(37), 11441–11450. doi:10.1523/JNEUROSCI.2387-09.2009

Clark, K., Sword, B. A., & Dupree, J. L. (2017). Oxidative Stress Induces Disruption of the Axon Initial Segment. ASN Neuro, 9(6), 1759091417745426. doi:10.1177/1759091417745426

Colledge, M., Snyder, E. M., Crozier, R. A., Soderling, J. A., Jin, Y., Langeberg, L. K., … Scott, J. D. (2003). Ubiquitination regulates PSD-95 degradation and AMPA receptor surface expression. Neuron, 40(3), 595–607. doi:10.1016/s0896-6273(03)00687-1

de Chaumont, F., Dallongeville, S., Chenouard, N., Herve, N., Pop, S., Provoost, T., … Olivo-Marin, J. C. (2012). Icy: an open bioimage informatics platform for extended reproducible research. Nat Methods, 9(7), 690–696. doi:10.1038/nmeth.2075

Del Puerto, A., Fronzaroli-Molinieres, L., Perez-Alvarez, M. J., Giraud, P., Carlier, E., Wandosell, F., … Garrido, J. J. (2015). ATP-P2X7 Receptor Modulates Axon Initial Segment Composition and Function in Physiological Conditions and Brain Injury. Cereb Cortex, 25(8), 2282–2294. doi:10.1093/cercor/bhu035

Dumitrescu, A. S., Evans, M. D., & Grubb, M. S. (2016). Evaluating Tools for Live Imaging of Structural Plasticity at the Axon Initial Segment. Front Cell Neurosci, 10, 268. doi:10.3389/fncel.2016.00268

Eichel, K., Uenaka, T., Belapurkar, V., Lu, R., Cheng, S., Pak, J. S., … Shen, K. (2022). Endocytosis in the axon initial segment maintains neuronal polarity. Nature. doi:10.1038/s41586-022-05074-5

Evans, M. D., Dumitrescu, A. S., Kruijssen, D. L. H., Taylor, S. E., & Grubb, M. S. (2015). Rapid Modulation of Axon Initial Segment Length Influences Repetitive Spike Firing. Cell Rep, 13(6), 1233–1245. doi:10.1016/j.celrep.2015.09.066

Evans, M. D., Sammons, R. P., Lebron, S., Dumitrescu, A. S., Watkins, T. B., Uebele, V. N., … Grubb, M. S. (2013). Calcineurin signaling mediates activity-dependent relocation of the axon initial segment. J Neurosci, 33(16), 6950–6963. doi:10.1523/JNEUROSCI.0277-13.2013

Evans, M. D., Tufo, C., Dumitrescu, A. S., & Grubb, M. S. (2017). Myosin II activity is required for structural plasticity at the axon initial segment. Eur J Neurosci, 46(2), 1751–1757. doi:10.1111/ejn.13597

Fache, M. P., Moussif, A., Fernandes, F., Giraud, P., Garrido, J. J., & Dargent, B. (2004). Endocytotic elimination and domain-selective tethering constitute a potential mechanism of protein segregation at the axonal initial segment. J Cell Biol, 166(4), 571–578. doi:10.1083/jcb.200312155

Ferreira da Silva, T., Granadeiro, L. S., Bessa-Neto, D., Luz, L. L., Safronov, B. V., & Brites, P. (2021). Plasmalogens regulate the AKT-ULK1 signaling pathway to control the position of the axon initial segment. Prog Neurobiol, 205, 102123. doi:10.1016/j.pneurobio.2021.102123

Freal, A., Rai, D., Tas, R. P., Pan, X., Katrukha, E. A., van de Willige, D., … Hoogenraad, C. C. (2019). Feedback-Driven Assembly of the Axon Initial Segment. Neuron. doi:10.1016/j.neuron.2019.07.029

Galliano, E., Hahn, C., Browne, L. P., P, R. V., Tufo, C., Crespo, A., & Grubb, M. S. (2021). Brief Sensory Deprivation Triggers Cell Type-Specific Structural and Functional Plasticity in Olfactory Bulb Neurons. J Neurosci, 41(10), 2135–2151. doi:10.1523/JNEUROSCI.1606-20.2020

Garrido, J. J., Giraud, P., Carlier, E., Fernandes, F., Moussif, A., Fache, M. P., … Dargent, B. (2003). A targeting motif involved in sodium channel clustering at the axonal initial segment. Science, 300(5628), 2091–2094. doi:10.1126/science.1085167

Gasser, A., Ho, T. S., Cheng, X., Chang, K. J., Waxman, S. G., Rasband, M. N., & Dib-Hajj, S. D. (2012). An ankyrinG-binding motif is necessary and sufficient for targeting Nav1.6 sodium channels to axon initial segments and nodes of Ranvier. J Neurosci, 32(21), 7232–7243. doi:10.1523/JNEUROSCI.5434-11.2012

Ghosh, A., Malavasi, E. L., Sherman, D. L., & Brophy, P. J. (2020). Neurofascin and Kv7.3 are delivered to somatic and axon terminal surface membranes en route to the axon initial segment. Elife, 9. doi:10.7554/eLife.60619

Grubb, M. S., & Burrone, J. (2010). Activity-dependent relocation of the axon initial segment fine-tunes neuronal excitability. Nature, 465(7301), 1070–1074. doi:10.1038/nature09160

Jamann, N., Dannehl, D., Lehmann, N., Wagener, R., Thielemann, C., Schultz, C., … Engelhardt, M. (2021). Sensory input drives rapid homeostatic scaling of the axon initial segment in mouse barrel cortex. Nat Commun, 12(1), 23. doi:10.1038/s41467-020-20232-x

Kamal, A., Ramakers, G. M., Urban, I. J., De Graan, P. N., & Gispen, W. H. (1999). Chemical LTD in the CA1 field of the hippocampus from young and mature rats. Eur J Neurosci, 11(10), 3512–3516. doi:10.1046/j.1460-9568.1999.00769.x

Kole, M. H., & Stuart, G. J. (2012). Signal processing in the axon initial segment. Neuron, 73(2), 235–247. doi:10.1016/j.neuron.2012.01.007

Kuba, H., Oichi, Y., & Ohmori, H. (2010). Presynaptic activity regulates Na(+) channel distribution at the axon initial segment. Nature, 465(7301), 1075–1078. doi:10.1038/nature09087

Kuba, H., Yamada, R., Ishiguro, G., & Adachi, R. (2015). Redistribution of Kv1 and Kv7 enhances neuronal excitability during structural axon initial segment plasticity. Nat Commun, 6, 8815. doi:10.1038/ncomms9815

Lee, H. K., Kameyama, K., Huganir, R. L., & Bear, M. F. (1998). NMDA induces long-term synaptic depression and dephosphorylation of the GluR1 subunit of AMPA receptors in hippocampus. Neuron, 21(5), 1151–1162. doi:10.1016/s0896-6273(00)80632-7

Lee, H. K., Takamiya, K., He, K., Song, L., & Huganir, R. L. (2010). Specific roles of AMPA receptor subunit GluR1 (GluA1) phosphorylation sites in regulating synaptic plasticity in the CA1 region of hippocampus. J Neurophysiol, 103(1), 479–489. doi:10.1152/jn.00835.2009

Lee, M., Liu, Y. C., Chen, C., Lu, C. H., Lu, S. T., Huang, T. N., … Cheng, P. L. (2020). Ecm29-mediated proteasomal distribution modulates excitatory GABA responses in the developing brain. J Cell Biol, 219(2). doi:10.1083/jcb.201903033

Lemaillet, G., Walker, B., & Lambert, S. (2003). Identification of a conserved ankyrin-binding motif in the family of sodium channel alpha subunits. J Biol Chem, 278(30), 27333–27339. doi:10.1074/jbc.M303327200

Leterrier, C. (2018). The Axon Initial Segment: An Updated Viewpoint. J Neurosci, 38(9), 2135–2145. doi:10.1523/JNEUROSCI.1922-17.2018

Leterrier, C., Clerc, N., Rueda-Boroni, F., Montersino, A., Dargent, B., & Castets, F. (2017). Ankyrin G Membrane Partners Drive the Establishment and Maintenance of the Axon Initial Segment. Front Cell Neurosci, 11, 6. doi:10.3389/fncel.2017.00006

Leyton-Puig, D., Isogai, T., Argenzio, E., van den Broek, B., Klarenbeek, J., Janssen, H., … Innocenti, M. (2017). Flat clathrin lattices are dynamic actin-controlled hubs for clathrin-mediated endocytosis and signalling of specific receptors. Nat Commun, 8, 16068. doi:10.1038/ncomms16068

Liu, H., Wang, H. G., Pitt, G. S., & Liu, Z. J. (2022). Direct Observation of Compartment-Specific Localization and Dynamics of Voltage-Gated Sodium Channels. J Neurosci. doi:10.1523/JNEUROSCI.0086-22.2022

Lorincz, A., & Nusser, Z. (2010). Molecular identity of dendritic voltage-gated sodium channels. Science, 328(5980), 906–909. doi:10.1126/science.1187958

Man, H. Y., Lin, J. W., Ju, W. H., Ahmadian, G., Liu, L., Becker, L. E., … Wang, Y. T. (2000). Regulation of AMPA receptor-mediated synaptic transmission by clathrin-dependent receptor internalization. Neuron, 25(3), 649–662. doi:10.1016/s0896-6273(00)81067-3

Oh, M. C., Derkach, V. A., Guire, E. S., & Soderling, T. R. (2006). Extrasynaptic membrane trafficking regulated by GluR1 serine 845 phosphorylation primes AMPA receptors for long-term potentiation. J Biol Chem, 281(2), 752–758. doi:10.1074/jbc.M509677200

Phair, R. D., Gorski, S. A., & Misteli, T. (2004). Measurement of dynamic protein binding to chromatin in vivo, using photobleaching microscopy. Methods Enzymol, 375, 393–414. doi:10.1016/s0076-6879(03)75025-3

Roth, R. H., Zhang, Y., & Huganir, R. L. (2017). Dynamic imaging of AMPA receptor trafficking in vitro and in vivo. Curr Opin Neurobiol, 45, 51–58. doi:10.1016/j.conb.2017.03.008

Schafer, D. P., Jha, S., Liu, F., Akella, T., McCullough, L. D., & Rasband, M. N. (2009). Disruption of the axon initial segment cytoskeleton is a new mechanism for neuronal injury. J Neurosci, 29(42), 13242–13254. doi:10.1523/JNEUROSCI.3376-09.2009

Schindelin, J., Arganda-Carreras, I., Frise, E., Kaynig, V., Longair, M., Pietzsch, T., … Cardona, A. (2012). Fiji: an open-source platform for biological-image analysis. Nat Methods, 9(7), 676–682. doi:10.1038/nmeth.2019

Sole, L., Wagnon, J. L., Akin, E. J., Meisler, M. H., & Tamkun, M. M. (2019). The MAP1B binding domain of Nav1.6 is required for stable expression at the axon initial segment. J Neurosci. doi:10.1523/JNEUROSCI.2771-18.2019

Willems, J., de Jong, A. P. H., Scheefhals, N., Mertens, E., Catsburg, L. A. E., Poorthuis, R. B., … MacGillavry, H. D. (2020). ORANGE: A CRISPR/Cas9-based genome editing toolbox for epitope tagging of endogenous proteins in neurons. PLoS Biol, 18(4), e3000665. doi:10.1371/journal.pbio.3000665

Winckler, B., Forscher, P., & Mellman, I. (1999). A diffusion barrier maintains distribution of membrane proteins in polarized neurons. Nature, 397(6721), 698–701. doi:10.1038/17806

Wong, E. H., Kemp, J. A., Priestley, T., Knight, A. R., Woodruff, G. N., & Iversen, L. L. (1986). The anticonvulsant MK-801 is a potent N-methyl-D-aspartate antagonist. Proc Natl Acad Sci U S A, 83(18), 7104–7108. doi:10.1073/pnas.83.18.7104

Wong, H. H., Rannio, S., Jones, V., Thomazeau, A., & Sjostrom, P. J. (2021). NMDA receptors in axons: there’s no coincidence. J Physiol, 599(2), 367–387. doi:10.1113/JP280059

Woodruff, G. N., Foster, A. C., Gill, R., Kemp, J. A., Wong, E. H., & Iversen, L. L. (1987). The interaction between MK-801 and receptors for N-methyl-D-aspartate: functional consequences. Neuropharmacology, 26(7B), 903–909. doi:10.1016/0028-3908(87)90068-2

Xu, J., Kang, N., Jiang, L., Nedergaard, M., & Kang, J. (2005). Activity-dependent long-term potentiation of intrinsic excitability in hippocampal CA1 pyramidal neurons. J Neurosci, 25(7), 1750–1760. doi:10.1523/JNEUROSCI.4217-04.2005

Yoon, S., Parnell, E., Kasherman, M., Forrest, M. P., Myczek, K., Premarathne, S., … Penzes, P. (2020). Usp9X Controls Ankyrin-Repeat Domain Protein Homeostasis during Dendritic Spine Development. Neuron, 105(3), 506–521 e507. doi:10.1016/j.neuron.2019.11.003

Zhang, W., & Linden, D. J. (2003). The other side of the engram: experience-driven changes in neuronal intrinsic excitability. Nat Rev Neurosci, 4(11), 885–900. doi:10.1038/nrn1248

Zhao, Y., Wu, X., Chen, X., Li, J., Tian, C., Chen, J., … He, S. (2020). Calcineurin Signaling Mediates Disruption of the Axon Initial Segment Cytoskeleton after Injury. iScience, 23(2), 100880. doi:10.1016/j.isci.2020.100880

