## Supplementary Material for "Sodium channel endocytosis drives axon initial segment plasticity"

**This PDF file includes:**

Figs. S1 to S7

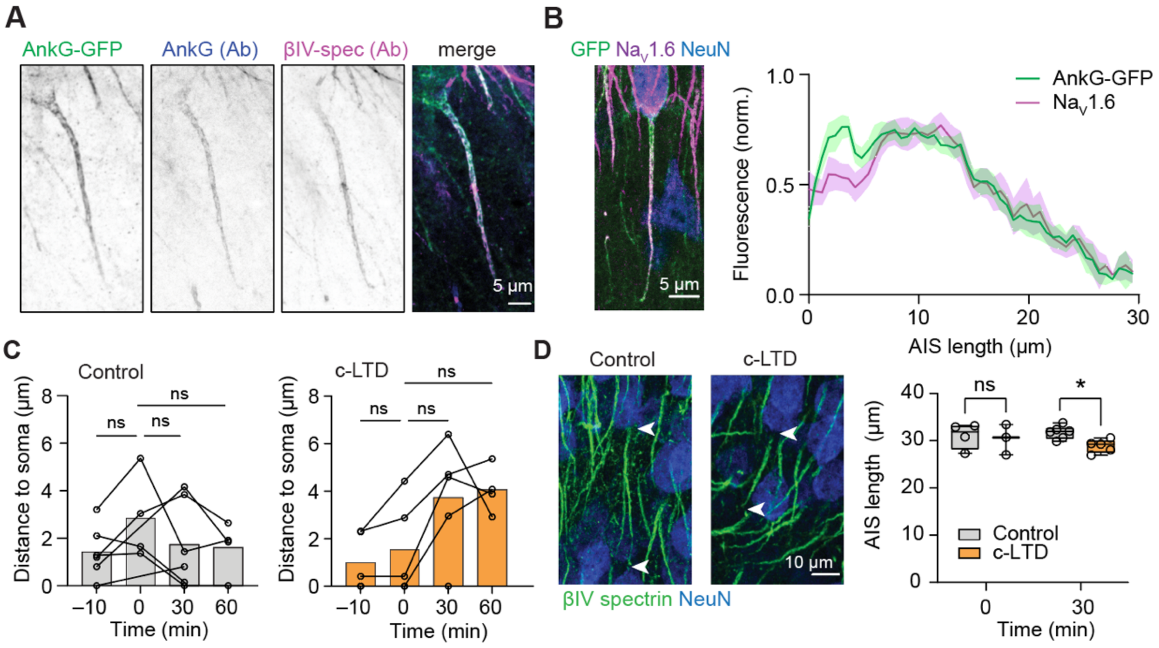


**Fig. S1. Validation of AIS targeting of AnkG-GFP and NMDAR-mediated plasticity**

(**A**) Example immunofluorescence GFP signal was localized to the AIS and overlapped with AnkG (gp Ab) as well as β4-spectrin (rb) antibody stainings (**B**) Fluorescence intensity distribution shows that Na$_V$1.6 (magenta) overlaps with the AnkG-GFP signals (green) along the AIS (*n* = 10 AIS). Lines show average (continuous) with SEM (transparent area). (**C**) Analysis of the distance of the AIS onset relative to the soma edge for control (*n* = 6 neurons, grey)) and c-LTD (*n* = 5 neurons, orange). Mixed-effects model (REML) p = 0.355 (control) and *p = 0.047 (c-LTD) for the factor treatment. Dunnett's multiple comparisons tests for all comparisons p > 0.05 (ns). (**D**) Incubation of acute hippocampal slices in 20 µM NMDA for 3 min followed by 5 min APV (c-LTD) vs APV treatment alone (5 min treatment, Control) reveals a significant shortening of the AIS, stained with β4-spectrin, after 30 min recovery. Scale bar is 10 µm Mixed-effects model *p = 0.017 for the factor treatment. Šídák's multiple comparisons tests ns p = 0.43 (0 min) and *p = 0.017 (30 min).

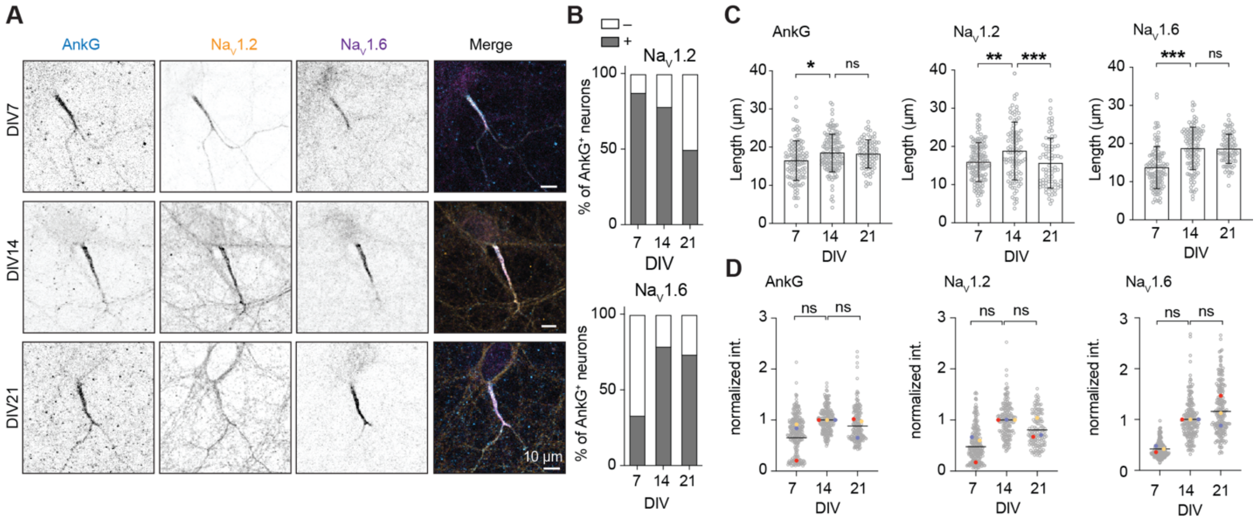


**Fig. S2. AIS expression of $Na_V1.2$ and $Na_V1.6$ during hippocampal neurons development in culture**

Immunostaining of AnkG, $Na_V1.2$ and $Na_V1.6$ in cultured hippocampal neurons at DIV7, DIV14 and DIV21. Scale bars are 10 µm. **(B)** Percentage of neurons with AnkG-positive AIS staining positive (grey) or negative (white) for $Na_V1.2$ or $Na_V1.6$. **(C)** Length of AnkG, $Na_V1.2$ or $Na_V1.6$ at DIV7, DIV14 and DIV21. Ordinary one-way ANOVA. $n$ = 77 to142 neurons (grey) from 3 experiments (colored dots). For AnkG, DIV14 **p = 0.0025, DIV21 ns p = 0.92. For $Na_V1.2$, DIV14 **p = 0.0013, DIV21 ***p = 0.0024. **(D)** Normalized AIS fluorescence intensity of AnkG, $Na_V1.2$ or $Na_V1.6$ staining at DIV7, DIV14 and DIV21. Kruskal-Wallis test with Dunn's multiple comparison test, $N$ = 3 experiments (22 to 124 neurons per condition per experiment). For AnkG, DIV14 ns p = 0.14, DIV21 ns p>0.99. For $Na_V1.2$, DIV14 ns p = 0.07, DIV21 ns p > 0.99. For $Na_V1.6$, DIV14 ns p = 0.20, DIV21 ns p > 0.99.

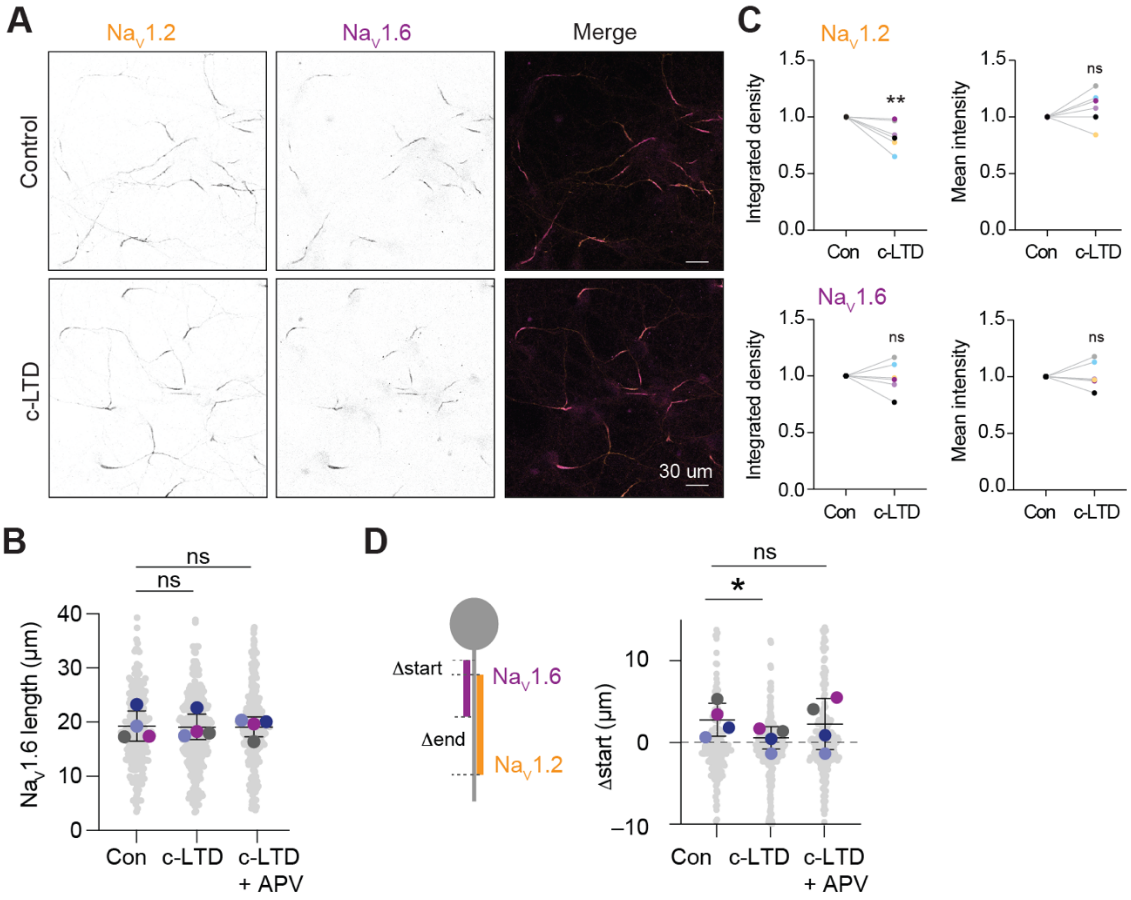


**Fig.S3 Na$_V$1.2 integrated density at the AIS is affected by NMDAR-mediated plasticity, whereas Na$_V$1.6 distribution remains unaltered**

**(A)** DIV14 hippocampal neurons stained for both Na$_V$1.2 and Na$_V$1.6, scale bars are 30 µm
(**B**) AIS length of Na$_V$1.6 in control, c-LTD and c-LTD+APV conditions was not affected (Repeated-measure one-way ANOVA with Dunnett's multiple comparisons test, $N = 4$ cultures. Control vs c-LTD, ns = 0.93; control vs c-LTD + APV, ns p = 0.98, >225 neurons per culture).
**(C)** Normalized integrated and mean intensity of Na$_V$1.2 and Na$_V$1.6 in control condition and after c-LTD. Unpaired t-test, N = 5 cultures (at least 30 neurons per condition per experiment). For Na$_V$1.2, integrated density **p = 0.009, mean intensity ns p = 0.19. For Na$_V$1.6, integrated density ns p = 0.79, mean intensity ns p = 0.79.
(**D**) Δstart (Na$_V$1.2 minus Na$_V$1.6 signal onset) along the AIS in control, c-LTD and c-LTD+APV conditions. RM ANOVA with Dunnett's multiple comparisons test, $N = 4$ cultures. c-LTD, *p = 0.048, c-LTD+APV, ns p = 0.80, >260 neurons per condition.

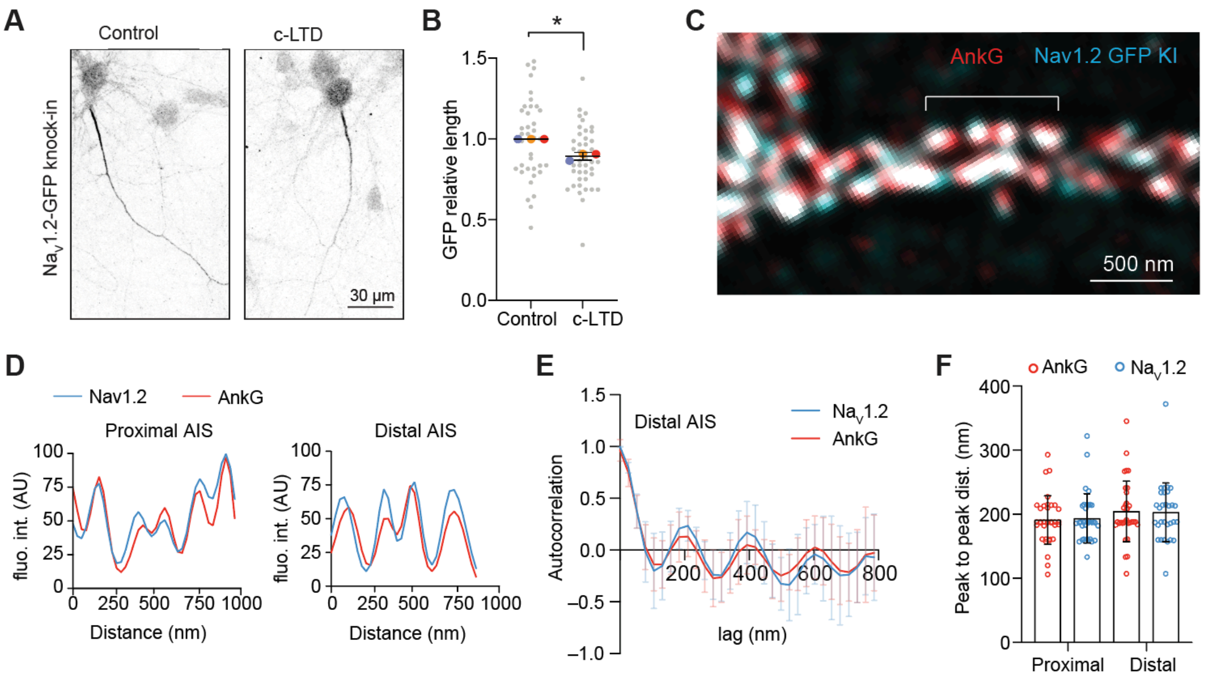


**Fig. S4. The knock-in of GFP in the C-terminal region of Na$_V$1.2 does not alter the nanoscale organization nor activity-dependent plasticity**

**(A,B)** Confocal images of DIV14 Na$_V$1.2-GFP knock-in neurons stained for GFP in control condition and after c-LTD. Na$_V$1.2-GFP length in c-LTD relative to the control condition, Unpaired t-test, N = 3 independent experiments (colored dots), **p = 0.0019 (*n* = 48 to 64 neurons, grey dots).
**(C)** STED image of the distal portion of the AIS in a DIV14 Na$_V$1.2-GFP knock-in neuron stained for GFP and AnkG. Scale bar is 500 nm. **(D)** Fluorescence intensity profile of AnkG and Na$_V$1.2 in the proximal and distal AIS, along the brackets shown in (C) and in main Figure 2 panel (F). (**E**) Autocorrelation profiles of AnkG and Na$_V$1.2 fluorescence intensity in distal AIS regions, *n* = 8 neurons. (**F**) Mean peak to peak distance of the autocorrelation profiles for AnkG and Na$_V$1.2 in the proximal and distal AIS from *n* = 30 to 34 measures from 8 neurons. Kruskal-Wallis test followed by a Dunn's multiple comparison test, p > 0.99 for all comparisons.

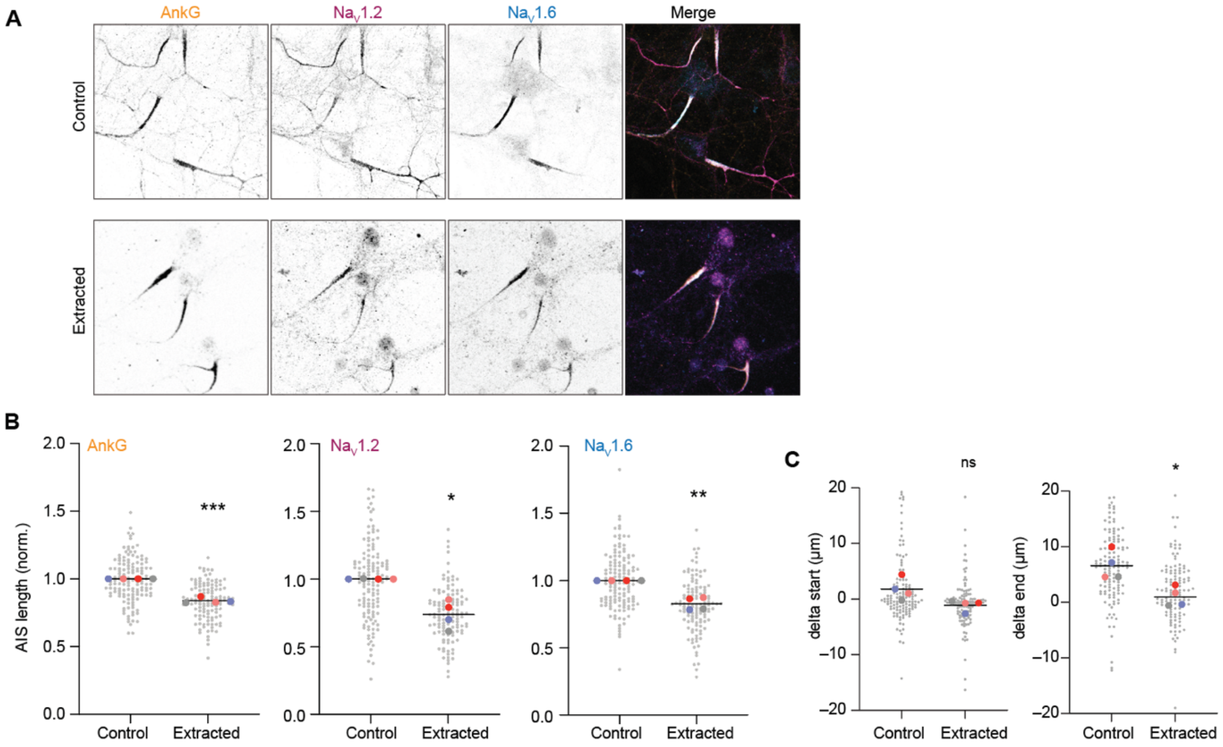


**Fig. S5. Resistance of Na$_V$1.2 and Na$_V$1.6 to triton extraction**

**(A)** DIV14 hippocampal neurons in control condition or after Triton extraction stained for AnkG, Na$_V$1.2 and Na$_V$1.6. Scale bars are 20 µm. **(B)** Length of AnkG, Na$_V$1.2 and Na$_V$1.6 after extraction relative to the control condition. *N* = 4 experiments, Unpaired t-test, for AnkG ***p < 0.0001, for Na$_V$1.2 **p = 0.002 and for Na$_V$1.6 ***p=0.0004, at least 65 neurons per condition per experiment. (**C**) Delta start and delta end positions (Na$_V$1.2 minus Na$_V$1.6 signal onset) along the AIS in control condition and after extraction. *N* = 4 experiments, Paired t-test, for delta start ns p = 0.08, for delta end *p = 0.012 (> 60 neurons per condition per experiment).

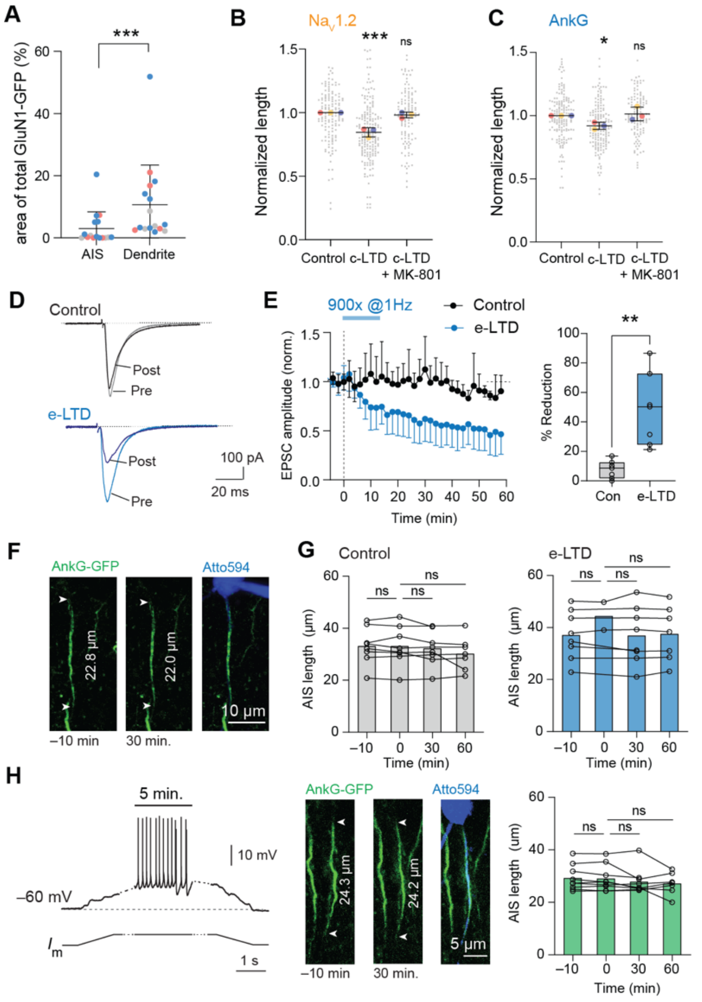


### Fig. S6. AIS stability during electrical LTD and action potential firing

**(A)** Analysis of total GFP staining area in the dendrites and AIS in neurons knock-in for GluN1-GFP. Wilcoxon test, *n* = 16 neurons, from 3 independent experiments ***p<0.0001.

(**B,C**) Length of $Na_V$1.2 (A) and AnkG (B) after c-LTD or c-LTD+MK-80 relative to control condition. N=3 experiments, Kruskal-Wallis test with Dunn's multiple comparison test. For AnkG c-LTD *p = 0.045, c-LTD+MK-801 ns p > 0.99. For $Na_V$1.2 c-LTD *p = 0.046, c-LTD+MK-801 ns p > 0.99. At least 80 neurons per condition per experiment.

(**D**) Example traces of SC PSC before (pre) and 60 min. after (post) electrical stimulation of Schaffer collaterals (900x @1 Hz, e-LTD). (**E**) e-LTD caused a significant reduction of PSC peak amplitude (*n* = 7) after 60 minutes of recording compared to controls (*n* = 7). Unpaired t-test **p = 0.0012. (**F**) Example confocal live-imaging of an AIS during e-LTD with stable AIS length. (**G**): No significant length changes. e-LTD (*n* = 8) vs control (*n* = 8) cells after 30 or 60 minutes of recording. Mixed-effects analysis with Dunnett's multiple comparisons test, p = 0.24 control, p = 0.43 e-LTD for the factor treatment. P > 0.05 for all posthoc multiple comparisons. (**H**) DC current injection produced neuronal action potential firing for ~5 minutes (*n* = 9) but a stable AIS length up to 60 minutes of recording and imaging. Mixed-effects analysis with Šídák's multiple comparisons test p = 0.118 for the factor treatment. P > 0.05 for all multiple comparisons.

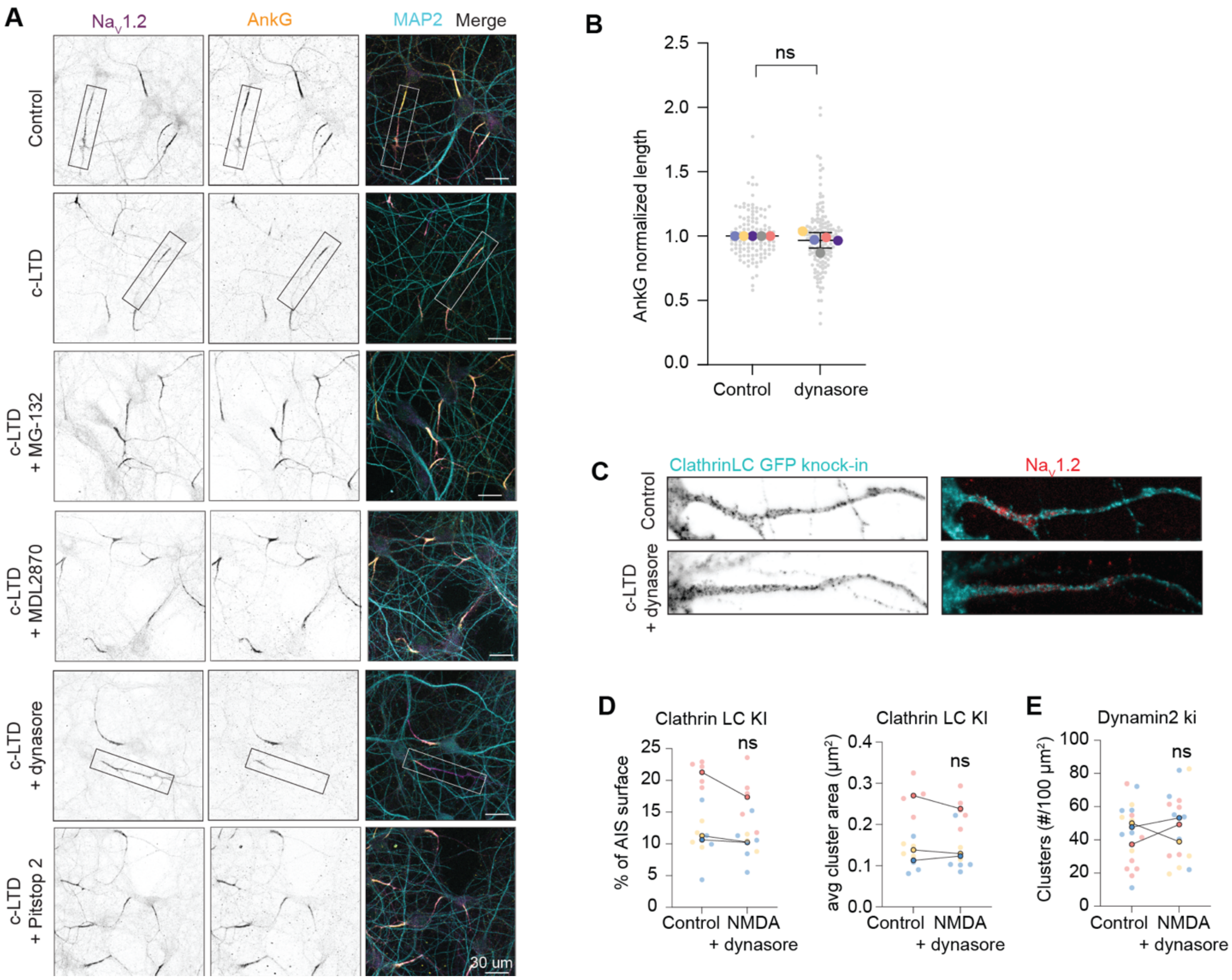


**Fig. S7. Blocking clathrin-mediated endocytosis prevents c-LTD induced AIS shortening**

(**A**) DIV14 hippocampal neurons in control condition, after c-LTD and in the presence of indicated drugs, stained for AnkG, $Na_V1.2$ and MAP2. Zooms of the boxed areas are shown in the main Figure 5, panel A. Scale bars are 20 µm.

(**B**) AnkG length after dynasore treatment relative to control condition. *N* = 5 cultures, unpaired t-test ns p = 0.26 (at least 30 neurons per condition per experiment).

(**C,D**) Confocal images of the AIS of ClathrinLC-GFP knock-in neurons stained for GFP and $Na_V1.2$ in control condition and after c-LTD in the presence of dynasore (C). Population analysis of clathrinLC mean cluster area and surface occupancy at the AIS (D). For the AIS occupancy: paired t-test, ns p = 0.24. For the cluster area: paired t-test, ns p = 0.49. *N* = 3 independent experiments, 4–5 neurons per condition per experiment.

(**E**) Density of endogenous dynamin2-GFP clusters. Paired t-test ns p = 0.78, *N* = 3 independent experiments, 4–7 neurons per condition per experiment.
